# Archetypal analysis provides interpretable mixture representations of microbiome community structure

**DOI:** 10.64898/2026.11.18.767757

**Authors:** Ray Verma, Ava Albert, Alexander G. Ioannidis, Aashish R. Jha

## Abstract

Microbiome studies routinely reduce high-dimensional community profiles to a small number of coordinates for visualization and comparison. Ordination methods are powerful for detecting compositional differences, but their axes are often difficult to translate into explicit microbial community configurations; conversely, hard clustering can impose discrete states on variation that is frequently continuous. Here we evaluate Archetypal Analysis (AA) as an intermediate representation in which each microbiome is expressed as a convex combination of a small number of data-supported extreme community profiles. Across three applications, AA recovered biologically recognizable structure while preserving graded variation among samples. In a multi-biospecimen human microbiome dataset, different sample-source categories occupied characteristic regions of archetype space, while stool samples were represented by heterogeneous mixtures of archetypal profiles. In an early-life gut microbiome dataset, archetype usage changed systematically with age, describing microbial succession as changing contributions from community endpoints. In domestic dogs spanning forager, agrarian, pastoralist, urban, and industrialized lifestyles, groups showed characteristic but non-discrete combinations of archetypes, supporting generality across host species and ecological contexts. Across the displayed K-sensitivity analyses, the principal sample-source and early-life organization remained recognizable as model resolution changed, while repeated initialization and resampling analyses recovered similar archetypal structure. Held-out samples projected onto fixed training-derived archetypes retained metadata-associated organization within each dataset. Bray-Curtis principal coordinates analysis recovered the same broad biological structure, indicating that the value of AA is interpretive rather than simply greater separation, while taxa prioritized by AA retained substantial predictive information relative to equally sized random taxon sets. Together, these results support AA as a complementary microbiome representation that connects low-dimensional sample coordinates directly to interpretable taxonomic profiles and provides a natural framework for describing ecological states, gradients, and transitions without hard cluster assignments.

**Author Summary:** Microbiome datasets contain hundreds to thousands of microbial features for every sample, making them difficult to visualize and interpret. Researchers commonly reduce these data to mathematical coordinates or divide samples into clusters. Both strategies are useful, but coordinates can be biologically abstract and clusters can make gradual ecological changes appear artificially discrete. We evaluate archetypal analysis as a complementary approach. Instead of assigning each microbiome to a single group, the method identifies a small number of extreme community profiles, called archetypes, and describes each sample as a mixture of those archetypes. We apply this framework to human microbiomes collected from different sample types, gut microbiome development during early childhood, and gut microbiomes of dogs living across different lifestyles. In each setting, the archetypes capture recognizable biological structure while preserving variation within groups and along continuous gradients. We also show that taxa emphasized by the archetypes retain substantial information about the biological patterns in the data. We therefore propose Archetypal Analysis as an interpretable way to summarize microbiome variation while keeping the resulting coordinates directly connected to microbial composition.

## Introduction

Microbial communities vary across anatomical sites, hosts, environments, populations, and life stages, yet much of this high-dimensional variation is organized along recurring ecological patterns (1). In the human microbiome, these patterns include strong differences among body habitats, developmental trajectories during early life, and shifts associated with diet, geography, subsistence, and industrialization (2-6). Large microbiome compendia are making these gradients increasingly visible across thousands of samples (7). A central computational challenge is therefore not simply to detect that communities differ, but to represent those differences in a reduced form that remains biologically interpretable.

Ordination methods such as principal coordinates analysis (PCoA), principal component analysis (PCA), and non-metric multidimensional scaling (NMDS) have become indispensable for visualizing microbiome variation (1). They efficiently reveal dominant directions or distances among samples, but the resulting coordinates are mathematical summaries: a position on a principal coordinate or component does not itself correspond to a microbial community profile. At the other conceptual extreme, clustering and community-typing approaches can provide intuitive group labels, but microbial variation is often continuous rather than cleanly partitioned.

Despite their utility, these methods often leave a gap between statistical representation and ecological interpretation. Distance-based embeddings such as PCoA preserve relationships between samples but do not directly identify representative community states. The history of gut enterotypes illustrates this tension: discrete community configurations can be useful summaries, yet the underlying landscape can contain dense regions connected by gradients rather than sharply separated states (8, 9). In practice, microbiome data are frequently interpreted in terms of ecological extremes and transitions, but most commonly used dimensionality-reduction approaches do not explicitly model such extremes or provide a framework for expressing individual samples as mixtures of biologically interpretable ecological states.

Archetypal Analysis (AA) offers a natural representation between these conceptual extremes. Introduced by Cutler and Breiman, AA approximates observations as convex combinations of a small number of archetypes, while constraining each archetype itself to be a convex combination of observed samples (10). Geometrically, AA seeks data-supported extremes of the multivariate distribution rather than orthogonal axes or average cluster centroids (10, 11). The resulting coefficients are continuous and sum to one, so each sample can be described by its relative affinity to several archetypal profiles. This provides explicit reference configurations without requiring hard assignment to a single state. Importantly, archetypes are defined in the original feature space and can therefore be interpreted directly as microbial community profiles. AA thus not only identifies the extreme configurations that organize community variation, but also the taxa that characterize each archetype. This formulation aligns naturally with an ecological view of microbiome variation, in which observed communities can be represented as combinations of extreme community configurations. AA has been developed for general machine learning and data mining (11, 12) and has been adapted to population genetics, where it provided interpretable ancestry proportions related to established clustering methods (13).

Interest in archetypal representations of microbiomes is also emerging. Recent work combined deep learning with AA to identify functional archetypes in nearly 10,000 adult gut metagenomes and linked those archetypes to microbiome stability and disease-associated signatures (14). A recent study used conventional AA to summarize activated-sludge microbiomes in relation to nitrous oxide emission states (15). These studies demonstrate the potential of archetypal representations, but a general and transparent evaluation of conventional AA as a practical representation of taxonomic microbiome structure across common study designs remains limited. In particular, it is not yet clear how AA behaves across strong habitat differences, continuous developmental succession, and lifestyle-associated variation, or how its outputs should be interpreted relative to conventional microbiome ordination.

Here, we evaluate AA as a general framework for representing microbiome community variation. Rather than asking whether AA replaces ordination, we ask whether it adds a complementary layer of interpretation: can known biological structure be expressed as mixtures of a small number of data-supported microbial extremes, and can those mixtures provide intuitive descriptions of both relatively discrete differences and continuous transitions? We examine a multi-biospecimen human microbiome dataset, an early-life gut microbiome dataset spanning infancy and childhood, and a domestic dog gut microbiome dataset spanning forager, agrarian, pastoralist, urban, and industrialized lifestyles. Across these examples, AA converts multivariate variation into archetypal community profiles and sample-specific mixture coefficients. We further examine sensitivity to the number of archetypes, reproducibility across initialization and data resampling, projection of held-out samples onto fixed archetypal references, and whether taxa prioritized by AA retain predictive information about sample metadata. Together, these analyses support AA as an interpretable coordinate system for microbiome states, gradients, and transitions.

## Results

### Archetypal Analysis represents microbiomes as mixtures of data-supported extreme profiles

For an abundance matrix X containing N microbiome samples and d microbial features, AA estimates K archetypal profiles and a K-component coefficient vector for each sample (**Fig. 1**). Each sample is approximated as a non-negative convex combination of the archetypes, and each archetype is constrained to lie within the convex hull of the observed data because it is itself a convex combination of samples (10, 11). The fitted sample coefficients therefore provide continuous coordinates that sum to one, while the archetypes can be mapped back to the original taxonomic feature space. This produces two directly linked views of the same data: where each sample lies relative to the archetypal extremes, and which taxa characterize those extremes.

**Figure 1.**
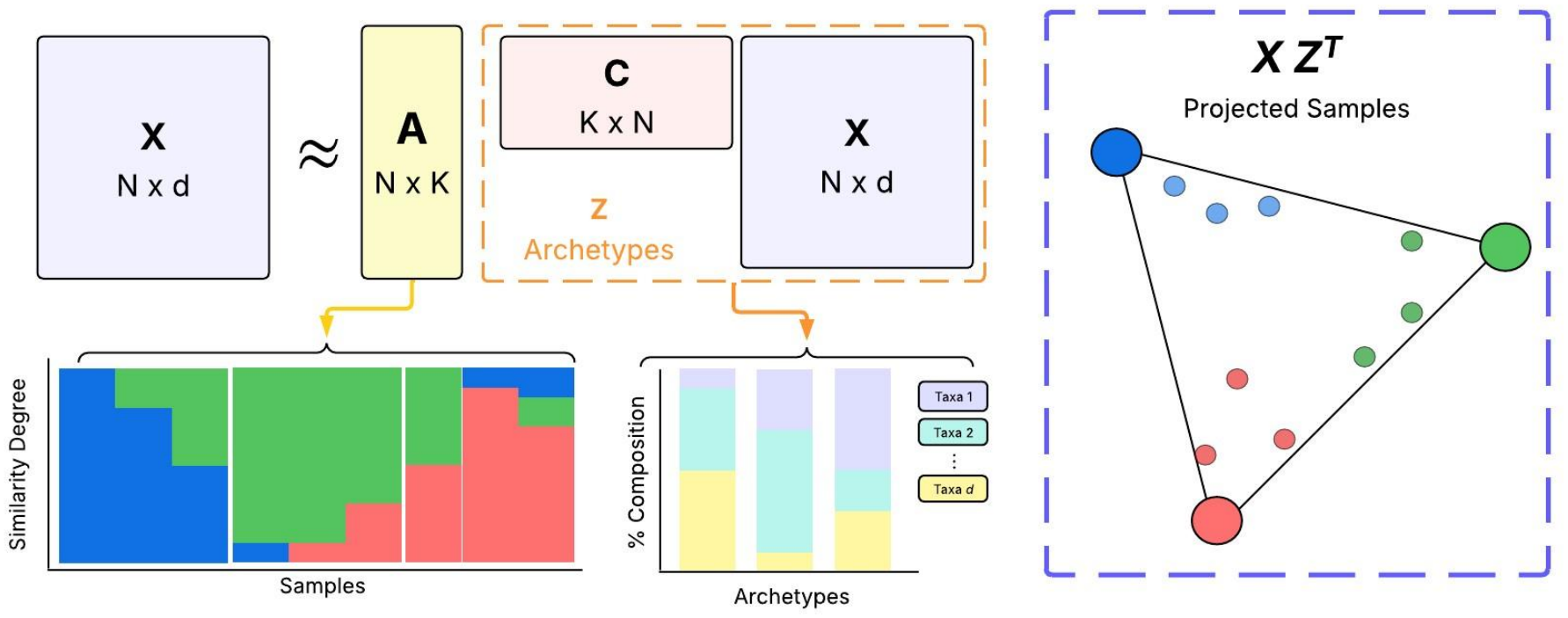
Archetypal Analysis framework for microbiome community profiles. The microbiome abundance matrix X is approximated by a sample-by-archetype mixing matrix and an archetype matrix derived from convex combinations of observed samples. Sample coefficients are non-negative and sum to one, allowing each microbiome to be represented as a mixture of K archetypes. Archetypes can be reconstructed in the original taxonomic feature space to identify the microbial features that characterize each extreme profile. A low-dimensional visualization of archetype coefficients provides an intuitive map of samples relative to the archetypal endpoints. The schematic is conceptual; for K greater than 3, any two-dimensional polygonal display is a projection of the K-component coefficient space rather than a literal K-simplex.

This representation differs conceptually from both ordination and clustering. PCoA and PCA summarize major dimensions of variation but do not define those coordinates as microbial community profiles; hard clustering assigns samples to groups but discards graded membership. AA instead describes each sample by its relative contributions from several data-supported endpoints. This mixture formulation is particularly useful for ecological transitions, where communities may move between endpoints rather than switch abruptly between discrete classes.

### Archetypal profiles recover established differences among human microbiome sample sources

We first asked whether AA could recover strong, well-established compositional differences among sample sources in a multi-biospecimen human microbiome dataset derived from the compendium reported by Abdill et al. (7). Seven sample-source categories were represented: appendiceal tissue (n = 58), cervicovaginal swab (n = 16), colorectal tumor (n = 56), esophageal biopsy (n = 44), fixed tissue (n = 35), ileal mucosal samples (n = 58), and stool (n = 52), for a total of 319 samples (**Fig. 2A**). We examined K = 3-10 and used K = 7 as a representative resolution for the primary display; sensitivity to model resolution and hard-assignment diagnostics are examined separately below (**Figs. S1-S2**). At K = 7, several sample-source categories concentrated near distinct archetypal regions. Appendiceal tissue was associated mainly with Archetypes 1 and 2, ileal mucosal samples with Archetype 3, colorectal tumor samples with Archetype 5, cervicovaginal swabs with Archetype 6, and esophageal biopsies with Archetype 7. In contrast, stool samples were distributed across several regions rather than concentrating near a single archetype. Because the stool samples were drawn from multiple studies, this broader dispersion cannot be attributed specifically to greater biological heterogeneity of stool microbiomes.

**Figure 2.**
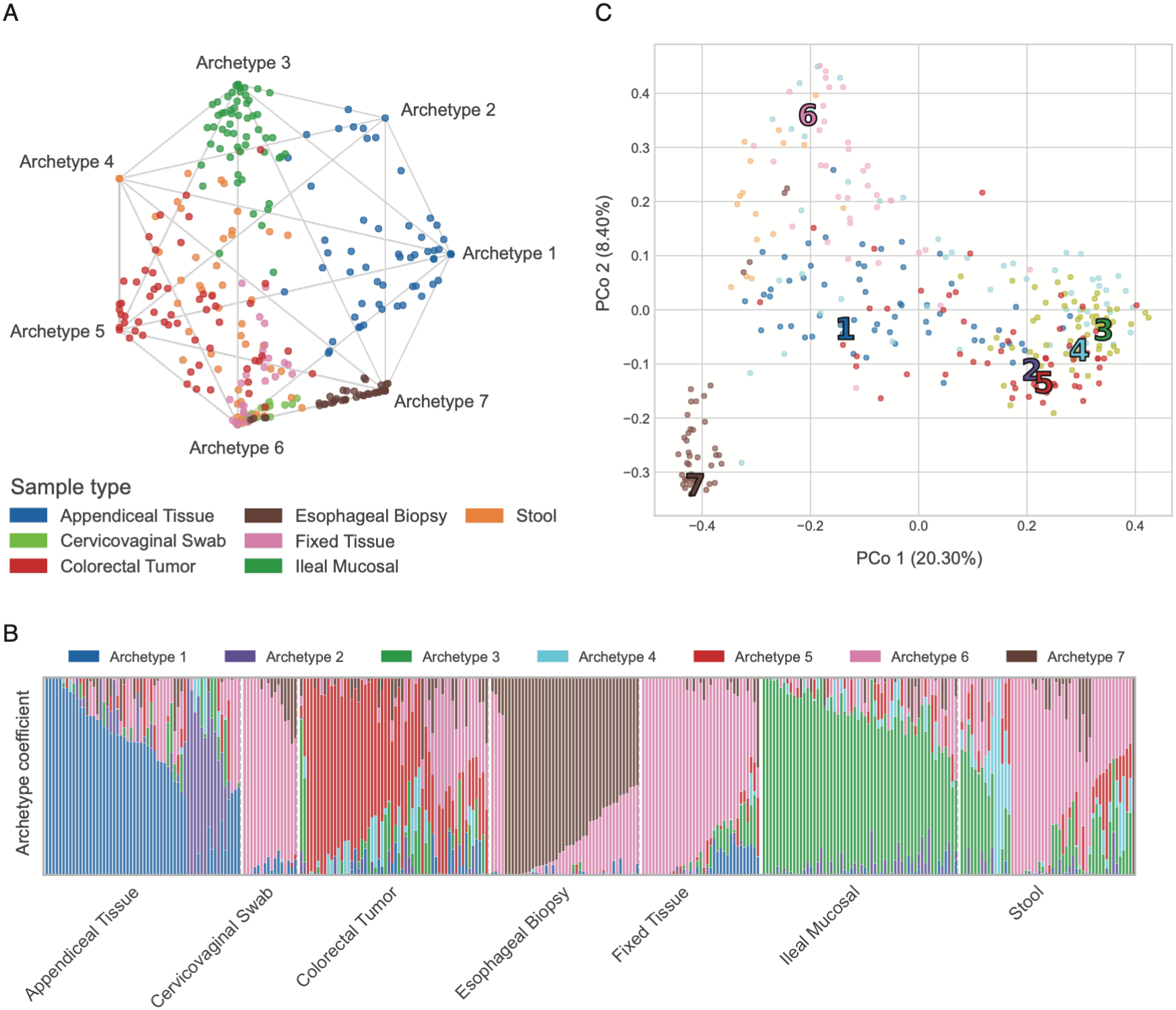
Archetypal Analysis organizes multi-biospecimen microbiome variation around sample-source-associated community extremes. **(A)** Two-dimensional visualization of the seven-component archetype coefficient space, with samples colored by sample-source category and archetypes positioned at the vertices of the polygonal projection. **(B)** Sample-wise archetype coefficients grouped by sample-source category. Several sample-source categories show strong affinity to particular archetype profiles, whereas others, including stool, are represented by mixtures of multiple archetypes. **(C)** Bray–Curtis principal coordinates analysis (PCoA) of the same samples, shown for PCo1 versus PCo2; numbered markers indicate the locations of the seven archetypal profiles projected into the sample ordination space.

The sample-level coefficient plot emphasized an important property of the representation: metadata categories were not reproduced as forced clusters. Categories with relatively distinctive community configurations showed high affinity to one or a few archetypes, whereas other categories retained mixed archetypal compositions (**Fig. 2B**). Thus, AA preserved within-category variation rather than requiring a one-to-one correspondence between sample-source labels and latent community states.

Bray-Curtis PCoA of the same samples also contained recognizable sample-source structure. PCo1 explained 20.30% of the variation, PCo2 8.40%, and PCo3 7.24% (**Fig. 2C, Fig. S3**). Esophageal biopsies separated strongly in PCo1-PCo2 space, whereas other sample-source differences became clearer across different combinations of principal coordinates. AA therefore did not reveal structure that was absent from conventional ordination. Instead, it provided an alternative representation of the broad compositional variation, organized around a common set of interpretable community endpoints. Importantly, these archetypes provide explicit taxonomic profiles of the inferred endpoints, allowing the taxa contributing to each extreme to be examined directly (**Fig. S4**).

Although Abdill et al. reprocessed the underlying sequence data using a standardized bioinformatic pipeline, the samples originated from multiple studies, and study of origin was confounded with sample-source category in this subset. Each non-stool sample-source category in the present subset was derived from a single BioProject, whereas the stool samples were drawn from 31 BioProjects. Consequently, sample-source effects cannot be separated from study-specific differences in sampling, sequencing protocol, disease context, or other upstream factors. We therefore use this dataset primarily as a positive-control demonstration that AA can recover and represent strong compositional structure, rather than to infer sample-source effects independent of study.

### Archetype mixtures provide an interpretable representation of early-life gut microbiome succession

We next examined a subset of the longitudinal gut microbiome data from The Environmental Determinants of Diabetes in the Young (TEDDY) study (5). The present analysis comprised 1,033 stool microbiome samples spanning approximately 2.2 to 72 months of age, all belonging to SRA study SRP154926 (BioProject PRJNA400115). These samples were processed using the standardized bioinformatics pipeline reported by Abdill et al. (7). Age was not used in fitting the archetypes. We examined K = 3-10 in the diagnostic analysis and used K = 6 as a representative resolution for the primary display; sensitivity to model resolution and hard-assignment diagnostics are examined separately below (**Figs. S5-S6**).

Despite being inferred without age information, the archetypal representation showed a clear developmental organization (**Fig. 3A**). The youngest samples were concentrated toward Archetype 1, whereas older samples increasingly occupied regions associated with other archetypes, including Archetype 6. Samples at intermediate ages showed contributions from multiple archetypes, producing a graded progression through archetype space rather than a series of discrete age-associated states.

**Figure 3.**
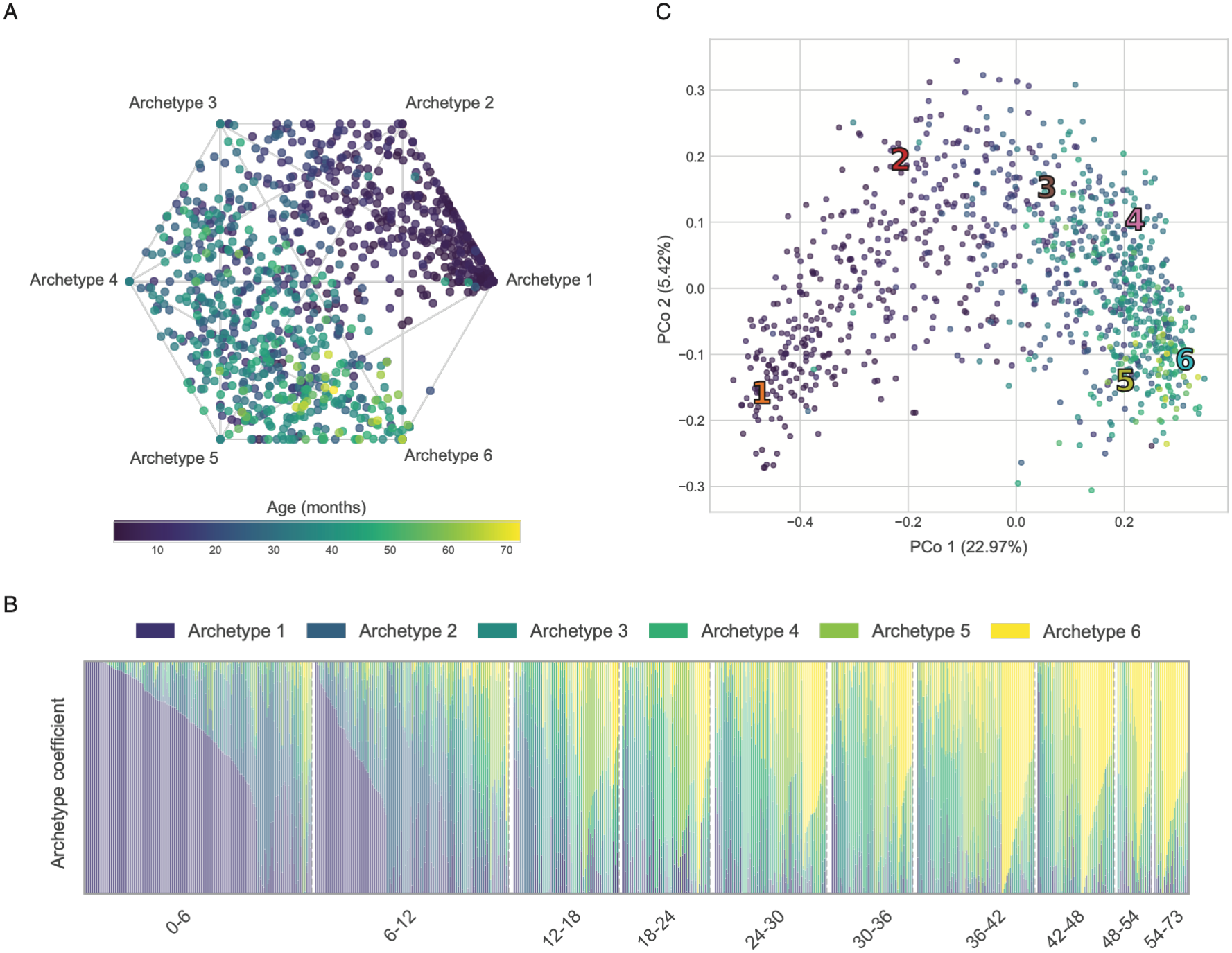
Archetype mixtures trace early-life gut microbiome succession. **(A)** Two-dimensional visualization of the six-component archetype coefficient space for the TEDDY early-life gut microbiome dataset, with samples colored continuously by age in months. **(B)** Sample-wise archetype coefficients grouped into successive age intervals from 0–6 to 54–73 months. Younger samples are dominated by Archetype 1, whereas contributions from other archetypes increase progressively with age, illustrating early-life microbiome succession as a graded shift among community profiles rather than a transition between discrete states. **(C)** Bray–Curtis principal coordinates analysis (PCoA) of the same samples, shown for PCo1 versus PCo2 and colored by age; numbered markers indicate the locations of the six archetypal profiles projected into the sample ordination space.

The sample-level coefficient representation further showed this developmental progression. Samples grouped across successive age intervals showed a decline in the dominance of the earliest-life archetype while alternative community configurations became progressively more prominent (**Fig. 3B**). Thus, microbiome development was not represented as a transition among a small number of discrete developmental states. Instead, different archetypal profiles contributed to different degrees across development, providing an interpretable representation of succession as changing mixtures of community endpoints.

Bray-Curtis PCoA of the same samples also captured a clear age-associated gradient. PCo1 explained 22.97% of the variation, PCo2 5.42%, and PCo3 3.93% (**Fig. 3C, Fig. S7**). AA therefore did not reveal developmental structure that was absent from conventional ordination. Rather, it represented the same biological progression in a form directly linked to explicit microbial community profiles. The archetypes showed distinct taxonomic compositions, including differences in *Bifidobacterium*, a hallmark genus of early-life gut microbiome development whose relative abundance changes with microbiome maturation (3, 5) (**Fig. S8**). Archetype 1 was also more compositionally concentrated than Archetype 6: its five highest-contributing taxa accounted for 52% of the archetypal profile, compared with 13% for Archetype 6. This developmental example therefore illustrates the interpretive value of the archetypal representation: continuous microbiome change can be described in terms of changing affinities to interpretable community endpoints rather than solely as movement along abstract ordination axes or transitions among discrete clusters.

### Lifestyle-associated dog microbiomes occupy characteristic mixtures of archetypal profiles

To test whether the framework generalized beyond humans and beyond strong anatomical or developmental differences, we applied AA to fecal microbiomes from domestic dogs sampled across forager, agrarian, pastoralist, urban, and industrialized lifestyles (16). The underlying cohort comprised 261 dogs from Nepal, Thailand, the United Arab Emirates, and the United States, with 257 retained after excluding recent antibiotic exposure (16). The original study identified lifestyle as the strongest measured correlate of canine gut microbiome composition, while also showing that lifestyle covaried with geography, climate, diet, and human management. We examined K = 3-10 and used K = 8 as a representative resolution for the primary display; sensitivity to model resolution and hard-assignment diagnostics are examined separately below (**Figs. S9-S10**).

With K = 8, the archetypal representation recapitulated this ecological structure while retaining substantial within-group heterogeneity (**Fig. 4A**). Industrialized dogs showed strong contributions from Archetypes 3 and 4, urban dogs were concentrated primarily toward Archetypes 1 and 2, pastoralist dogs were strongly associated with Archetype 5, and agrarian dogs showed contributions from Archetypes 6 and 8 (**Fig. 4B**). Forager dogs occupied a characteristic region of archetype space but also showed heterogeneous mixture coefficients. These patterns were visible in both the archetype map and the sample-level coefficient plot, where lifestyle categories displayed distinct distributions of mixture weights rather than one-to-one assignments to discrete states.

**Figure 4.**
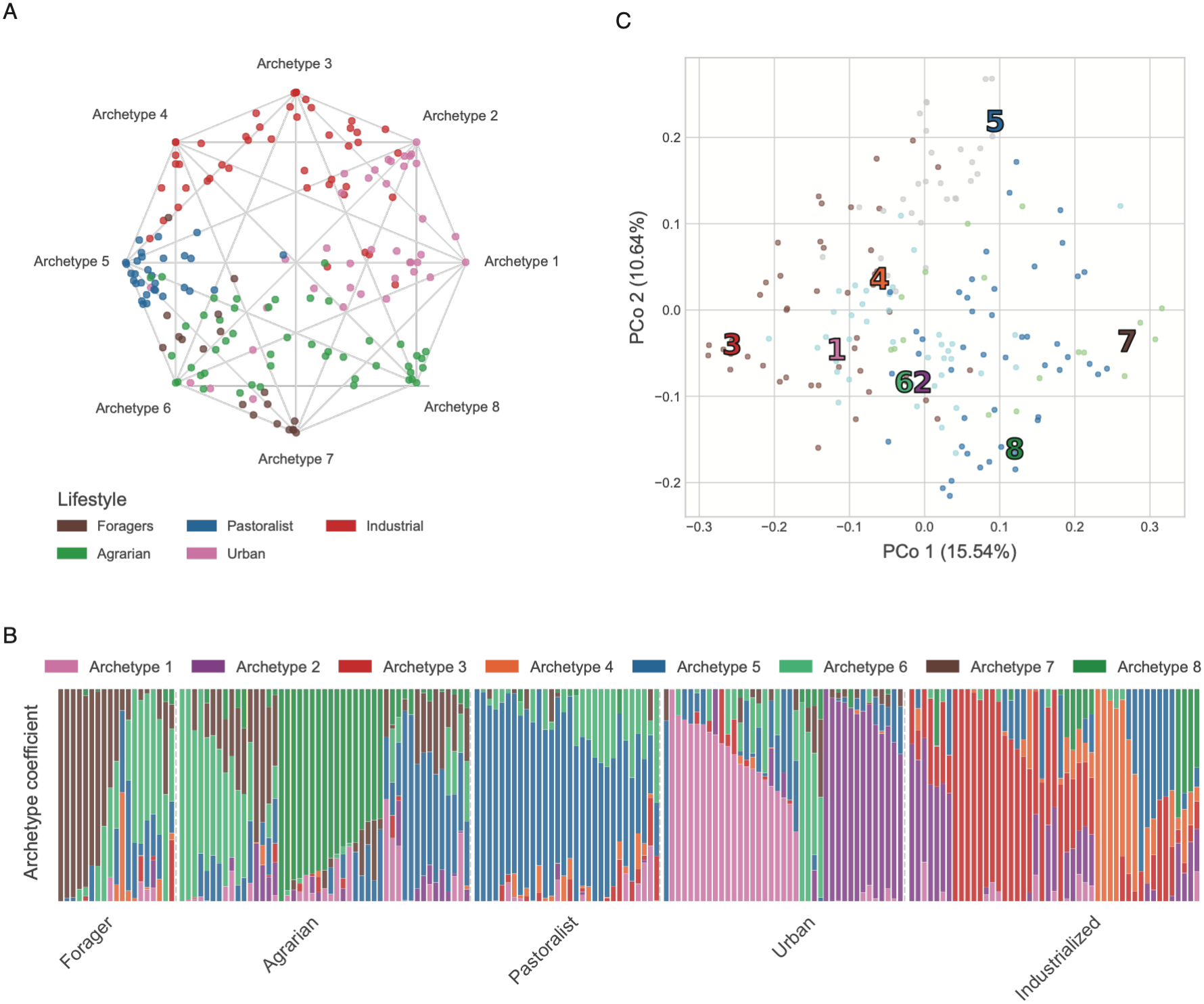
Dog gut microbiomes from different lifestyles occupy characteristic mixtures of archetypal profiles. **(A)** Two-dimensional visualization of the eight-component archetype coefficient space, with samples colored by lifestyle category. **(B)** Sample-wise archetype coefficients grouped by lifestyle. Lifestyle categories show characteristic distributions of archetype usage while retaining substantial within-group variation, illustrating that the archetypes represent continuous ecological endpoints rather than discrete lifestyle classes. **(C)** Bray–Curtis principal coordinates analysis (PCoA) of the same samples, shown for PCo1 versus PCo2; numbered markers indicate the locations of the eight archetypal profiles projected into the sample ordination space.

Bray-Curtis PCoA of the same samples also detected substantial lifestyle-associated structure. PCo1, PCo2, and PCo3 explained 15.54%, 10.64%, and 9.66% of the variation, respectively (**Fig. 4C, Fig. S11**). Industrialized, pastoralist, urban, and agrarian samples occupied recognizable regions across the first three coordinates. As in the other datasets, AA therefore did not reveal biological structure that was absent from conventional ordination. Rather, it provided an alternative representation of the same broad ecological variation in terms of interpretable community endpoints and their sample-specific mixtures. These profiles also identify the taxa characterizing the inferred ecological endpoints, providing a direct taxonomic interpretation of the archetypal variation observed across lifestyles **(Fig. S12)**.

Lifestyle in this cohort covaries with geography, climate, diet, age, and human management, and the observed archetype associations should therefore not be interpreted as causal effects of lifestyle itself. Instead, this dataset provides a test of whether AA can represent ecological variation within a single host species while preserving substantial variation within lifestyle groups.

### Taxa prioritized by archetypes retain predictive information about sample metadata

We next asked whether the microbial features emphasized by the archetypes were informative outside the AA fitting procedure. For each dataset, taxa were ranked according to their contributions to the archetypal profiles, and progressively larger sets of AA-prioritized taxa were used to train Random Forest models for the corresponding sample metadata. Performance was compared with equally sized randomly selected taxon sets (**Fig. 5**). Classification performance for sample source and dog lifestyle was evaluated using macro-F1, whereas prediction of age in the early-life dataset was evaluated using R^2^.

**Figure 5.**
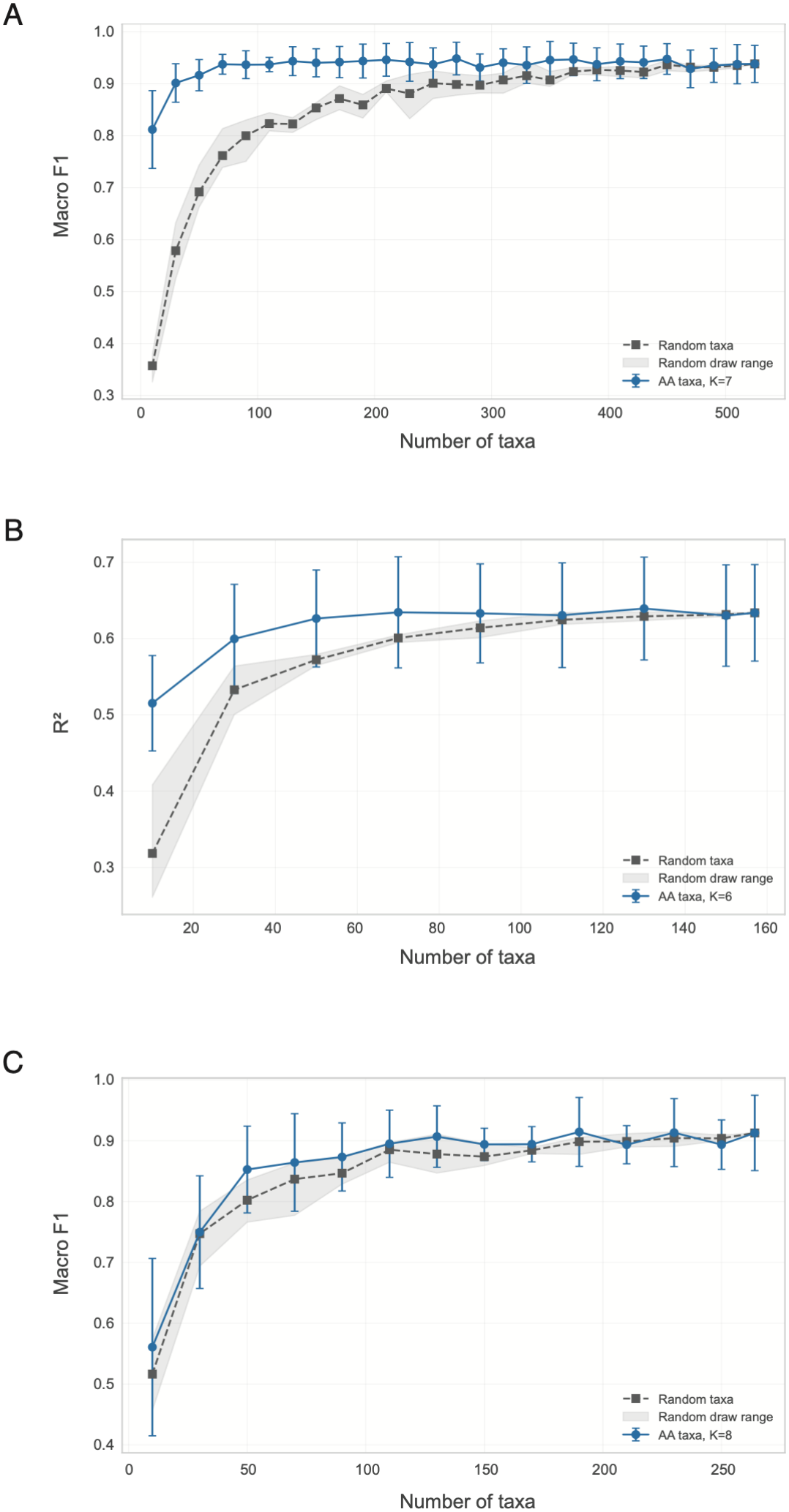
Taxa prioritized by archetypal profiles retain predictive information about sample metadata. Random Forest models were trained using progressively larger sets of taxa prioritized according to their contributions to the archetypal profiles and compared with models trained on equally sized randomly selected taxon sets. **(A)** Classification of human sample-source category using taxa prioritized from the K=7 archetypal representation; performance is shown as macro-F1. **(B)** Prediction of age in the TEDDY early-life dataset using taxa prioritized from the K=6 archetypal representation; performance is shown as R^2^. **(C)** Classification of dog lifestyle category using taxa prioritized from the K=8 archetypal representation; performance is shown as macro-F1. Blue points show performance using AA-prioritized taxa, black squares show performance using randomly selected taxa, and gray shading indicates variability across random taxon selections. Error bars show one standard deviation across model-evaluation replicates. The advantage of AA-prioritized taxa is greatest at smaller feature-set sizes and generally diminishes as increasing numbers of taxa are included.

For sample-source prediction, relatively small sets of AA-prioritized taxa achieved high macro-F1 values and substantially outperformed equally sized randomly selected feature sets. This advantage progressively diminished as larger numbers of taxa were included. In the early-life dataset, AA-prioritized taxa similarly retained substantial information about age, with particularly strong gains over random feature sets when relatively few taxa were used and convergence as feature-set size increased. In dogs, AA-prioritized taxa generally matched or modestly exceeded randomly selected taxa across much of the intermediate feature range, although the distinction was substantially weaker than for sample source or age.

This analysis is not intended to establish AA as a superior predictive model. Rather, it provides an independent test of interpretability: taxa contributing strongly to the archetypal endpoints concentrate information about the metadata-associated organization of the samples. The result complements the ordination analyses by linking the geometric extremes recovered by AA to compact sets of informative microbial features.

### Choice of K is treated as a resolution parameter rather than a hard clustering optimum

We evaluated archetypal analyses across multiple values of K and used silhouette and Davies-Bouldin statistics as secondary diagnostics after provisionally assigning each sample to the archetype with its largest mixture coefficient. The supplementary diagnostic plots cover K = 3-10 for all three datasets (**Figs. S1, S5, and S9**). Biological metadata were not used to fit AA. Because the primary AA representation is continuous, these temporary hard-assignment statistics were not treated as criteria defining a uniquely optimal number of archetypes.

The primary representations used K = 7 for sample source, K = 6 for early-life development, and K = 8 for dog lifestyle. These values were treated as parsimonious representative resolutions chosen for interpretability of the resulting mixture representation, while cross-K displays were used to assess whether the principal biological organization persisted as model resolution changed. The selected K values are therefore not presented as estimates of a discrete biological number of community states.

The diagnostic curves illustrate why this distinction matters. In the multi-biospecimen analysis, both the highest silhouette score and lowest Davies-Bouldin index within the examined range occurred at K = 10 (**Fig. S1**). Nevertheless, the broad sample-source organization was already apparent at lower resolutions and remained recognizable as additional archetypes refined the representation (**Fig. S2**); K = 7 was therefore used as a more parsimonious primary display. In TEDDY, silhouette coefficients remained slightly negative across K = 3-10 and Davies-Bouldin values varied across the examined range, with neither statistic identifying K = 6 as a unique optimum (**Fig. S5**). The age-associated organization nevertheless remained recognizable across resolutions (**Fig. S6**), and K = 6 was used as the primary representative resolution. In dogs, silhouette scores were relatively high and Davies-Bouldin values relatively low across approximately K = 8-10, without uniquely selecting K = 8 (**Fig. S9**). K = 8 was retained as the lowest representative resolution in this favorable range, with the corresponding cross-K organization shown in **Fig. S10**.

We therefore interpret K as a resolution parameter rather than as a hard clustering optimum. Increasing K can subdivide broad structure into additional archetypal endpoints, whereas lower values provide coarser summaries. Accordingly, our biological interpretations focus on organization that remains recognizable across reasonable changes in resolution rather than on the identity of individual archetypes at one supposedly optimal value of K; comparisons of specific archetypes across K require care because the fitted endpoints themselves can change.

### Archetypal profiles are reproducible across initialization and data resampling

To assess whether fitted archetypes depended strongly on optimization initialization or the particular samples used for fitting, we repeated AA across five random initializations of the full datasets and across repeated training subsets. Archetypes were matched between runs after accounting for label permutation, and stability was summarized using normalized Wasserstein distance. The three datasets yielded generally small distances across initializations; the distributions are shown separately rather than interpreted as exact invariance of the endpoints (**Fig. S13A**).

We next refit models to training subsets comprising 50%, 60%, 70%, 80%, or 90% of samples, with five random resampling replicates at each training fraction. Matched archetypal profiles remained broadly similar over these subsamples, although distances varied among datasets and across individual replicates (**Fig. S13B**). Together, the analyses support reproducibility of the broad fitted profiles under the perturbations examined, without establishing transferability to independent cohorts.

### Held-out samples can be projected onto fixed archetypal reference profiles

We next tested whether archetypes learned from a training subset could serve as a fixed reference for held-out samples. For each split, archetypes were fit using training samples only and were then held fixed while mixture coefficients were estimated for withheld samples. Across the plotted range of training fractions (10%-90%), the held-out projections preserved metadata-associated organization to varying degrees in all three datasets (**Fig. S14**). Silhouette, Davies-Bouldin, and PERMANOVA pseudo-F statistics summarize that organization in the held-out coefficient space; their interpretation depends on how metadata groups were defined, particularly for continuous age. These results demonstrate within-dataset projection to training-derived references, not validation of transportability across independent cohorts.

## Discussion

The central value of Archetypal Analysis for microbiome research is not that it reveals biological structure that ordination cannot detect. In all three applications examined here, conventional Bray-Curtis PCoA captured substantial aspects of the same ecological variation. Instead, AA provides a different representation of that variation. By anchoring coordinates to extreme microbial community profiles, it allows each microbiome to be interpreted as a graded combination of explicit biological configurations rather than only as a location along abstract axes or a member of a discrete cluster. Importantly, these archetypal profiles can be reconstructed in the original taxonomic feature space, directly identifying the microbial taxa that characterize each endpoint. AA therefore connects the geometry of community variation to the taxonomic composition of its underlying extremes, providing an interpretation that is not directly supplied by conventional ordination coordinates.

This distinction is most obvious when the biology itself is continuous. In the early-life dataset, age-related succession appears in PCoA as movement through multivariate coordinate space. AA describes the same developmental process in terms of changing contributions from archetypal microbial profiles. That representation is closer to the ecological questions researchers often want to ask: which community configurations characterize different stages of development, which configurations dominate intermediate stages, and how community composition shifts among them. Similar logic could be useful for perturbation and recovery, seasonal cycling, diet interventions, disease progression, and ecological restoration, all of which involve trajectories rather than simple case-control separation (2-6).

The sample-source and dog-lifestyle examples illustrate complementary properties. Strong biospecimen differences provide a positive control: unsupervised archetypes recover recognizable ecological endpoints, yet heterogeneous samples such as stool remain mixtures rather than being forced into a single state. The dog dataset demonstrates generality across host species and ecological variation within a single host species. Importantly, lifestyle groups do not map one-to-one onto archetypes. Industrialized, agrarian, pastoralist, urban, and forager dogs each show characteristic distributions of archetype usage, but substantial variation remains within groups. This is the behavior expected of a method intended to represent ecological continua rather than reproduce metadata labels.

The Random Forest analysis provides a complementary form of validation. Relatively small sets of taxa prioritized by the archetypal profiles retained substantial information about sample source, developmental age, and dog lifestyle and often outperformed equally sized sets of randomly selected taxa. This does not imply that AA-prioritized taxa are uniquely causal or that AA should be used as a supervised feature-selection algorithm. Rather, it shows that the microbial features defining the inferred extremes concentrate information about the metadata-associated organization of the samples, strengthening the interpretation of archetypes as meaningful summaries of community variation.

AA therefore occupies a useful conceptual space between ordination and clustering. It shares with ordination the ability to represent samples continuously and with clustering the ability to define interpretable reference profiles. The convex-hull constraint is important: archetypes are not unconstrained latent directions, but combinations of observations and therefore remain within the empirical domain of the dataset (10, 11). At the same time, archetypes should not be described as “pure” biological states or assumed to correspond to single observed samples. They are model-derived extremes whose biological meaning must be established through stability, taxonomic composition, biological associations, sensitivity to model resolution, and ultimately replication.

Conventional cluster-validity statistics reinforce the distinction between AA and hard clustering. Across the three datasets, silhouette and Davies-Bouldin curves did not consistently favor the representative K values used in the main figures. This is not unexpected because both metrics are defined on temporary dominant-archetype assignments rather than on the continuous mixture coefficients that constitute the primary AA output. In TEDDY, silhouette coefficients were negative throughout the plotted range, indicating weak separation under those temporary hard assignments; this observation does not by itself establish a biological continuum. For AA, K is better treated as a resolution parameter whose usefulness is judged by interpretability, stability, and the persistence of substantive conclusions across reasonable changes in resolution rather than by any single clustering index.

The initialization, resampling, and held-out analyses provide complementary robustness tests. Repeated fits recovered comparable archetypal structure across random initializations and partial training sets, and withheld samples could be assigned mixture coefficients relative to fixed training-derived archetypes while retaining the major metadata-associated organization within the same datasets. These results support the use of archetypes as a reproducible descriptive representation and indicate that training-derived archetypal profiles can provide a common coordinate system for samples not used during model fitting, while not establishing transportability across independent cohorts.

This framing also distinguishes the present study from recent archetypal microbiome work. Meawad et al. used a deep-learning extension of AA to characterize functional states of the adult human gut microbiome and demonstrated associations with stability and disease signatures (14). Chen et al. applied taxonomic AA to activated-sludge communities and linked archetypal profiles to nitrous oxide emission states (15). Our contribution is more methodological and general-purpose: to establish a transparent workflow for conventional AA on taxonomic microbiome profiles and to show how the same representation can describe habitat differences, developmental succession, and lifestyle-associated variation across host species. The practical value of this framework depends on reproducible implementation, robustness diagnostics, and clear guidance about what an archetype does and does not represent.

Several limitations require explicit consideration. First, archetypes are sampling-dependent. If biologically important extremes are absent or sparsely sampled, the inferred archetypes will change. Second, AA is sensitive by design to boundary observations, making resampling and stability assessment important for evaluating whether inferred archetypes depend strongly on particular samples. Third, the number of archetypes, K, is a modeling decision rather than a biological truth, and major interpretations should therefore be robust across reasonable changes in model resolution.

Fourth, microbiome abundance profiles are compositional (17), whereas the simplex constraint on AA coefficients is a separate mathematical property; AA should not be described as automatically solving compositional-data problems, and preprocessing choices should be explicitly reported and considered when interpreting results. Fifth, two-dimensional polygonal displays of K-component coefficients are visual projections when K exceeds three and should not be interpreted as literal geometries. Finally, observational metadata such as lifestyle can bundle geography, diet, climate, host characteristics, and technical factors; archetype associations should therefore be interpreted with the same causal restraint as conventional microbiome associations.

A further practical opportunity is to use archetypes as a fixed reference system. Once a set of archetypal taxonomic profiles has been learned in a training dataset, new samples can be assigned mixture coefficients relative to those fixed profiles without refitting the archetypes. The held-out projection analysis provides a direct test of this use case within the datasets examined here. Future evaluation in independent cohorts will be necessary to determine how well such reference systems transfer across studies, populations, and technical settings. If sufficiently stable, such a framework could allow longitudinal follow-up samples, external cohorts, or future observations to be placed in a common interpretable coordinate system. Archetype coefficients could then serve as low-dimensional phenotypes for association models, developmental trajectories, perturbation-recovery studies, or heterogeneous clinical groups. A lightweight implementation that accepts standard abundance tables or microbiome objects, reports K-sensitivity and stability diagnostics, reconstructs taxonomic archetypes, and supports held-out projection would make the method immediately useful to the broader microbiome community.

In summary, Archetypal Analysis offers a complementary way to represent microbiome community structure as extremes, mixtures, and transitions. The approach does not require the field to abandon established ordination methods. Rather, it adds a biologically grounded coordinate system in which dimensions correspond to microbial community profiles and individual samples can be interpreted by their graded affinities to those profiles. The present examples suggest that this representation is useful across anatomical, developmental, and ecological settings and warrants broader evaluation as a practical tool for microbiome analysis.

## Materials and Methods

### Overview of Archetypal Analysis

Archetypal Analysis was formulated following Cutler and Breiman (10). Let X be an N × d matrix of microbiome profiles, where N is the number of samples and d is the number of microbial features after preprocessing. For a chosen number of archetypes K, AA estimates a sample coefficient matrix A (N × K) and an archetype-definition matrix C (K × N). The K archetypal profiles are Z = CX, and the observed data are approximated as X ≈ AZ = ACX. AA minimizes the squared Frobenius reconstruction error ||X – ACX||^2^_F_ subject to convexity constraints on both coefficient matrices. Each row of A is non-negative and sums to one, so that each sample is represented as a convex combination of the K archetypes. Each row of C is likewise non-negative and sums to one, so that each archetype is itself a convex combination of observed samples (10, 11).

Throughout the manuscript, the coefficients in A are referred to as archetype usage, affinity, or mixture coefficients. These terms describe the geometric representation learned from the microbial abundance matrix and do not imply a literal ecological mixing mechanism. Archetypal profiles were reconstructed in the original microbial feature space to identify taxa contributing to each endpoint.

### Optimization and implementation

Archetypal Analysis was fit using alternating constrained optimization of the sample coefficients and archetype definitions, with non-negativity and sum-to-one constraints enforced at each step (10-12). The implementation developed for this study was used for all analyses. Primary representations used K = 7 for the human sample-source dataset, K = 6 for the early-life dataset, and K = 8 for the dog-lifestyle dataset.

All experiments were conducted using Python 3.11 and the archetypes package (18). Optimization was performed via Projected Gradient Descent (PGD) with AA++ initialization (19), employing a residual sum of squares (RSS) convergence tolerance of 10^−4^ and a maximum of 10,000 iterations.

### Microbiome datasets and preprocessing

#### General preprocessing

Analyses began from taxonomic count tables and associated sample metadata obtained from the corresponding public datasets or processed resources described below. Taxa lacking identification at the phylum level were removed, and profiles were analyzed at the genus level. Samples lacking required metadata or containing fewer than 10% of the taxa detected in the overall dataset, or having fewer than 5,000 total counts were excluded from analysis.

Taxonomic counts were aggregated at the genus level and log-transformed after addition of a pseudocount of 1 before PCoA and Archetypal Analysis. This transformation reduces the influence of highly abundant taxa and avoids information loss associated with rarefaction (20).

#### Human multi-biospecimen dataset

The sample-source analysis used microbiome profiles from the Human Microbiome Compendium reported by Abdill et al. (7). The analyzed subset comprised 319 samples from seven biospecimen categories: appendiceal tissue (n = 58), cervicovaginal swab (n = 16), colorectal tumor (n = 56), esophageal biopsy (n = 44), fixed tissue (n = 35), ileal mucosal samples (n = 58), and stool (n = 52). The primary Archetypal Analysis used K = 7. The exact dataset used for this analysis and its associated metadata are provided as a phyloseq object in Supplementary Dataset 1.

#### Early-life gut microbiome dataset

The early-life analysis used longitudinal stool microbiome data from The Environmental Determinants of Diabetes in the Young (TEDDY) study (5). The analyzed dataset comprised 1,033 samples spanning approximately 2.2 to 72 months of age from SRA study SRP154926 (BioProject PRJNA400115). Taxonomic profiles were obtained from the standardized processed microbiome resource reported by Abdill et al. (7). The primary Archetypal Analysis used K = 6. Age was not used during AA fitting and was introduced only for post hoc interpretation of the microbiome-derived representation. The exact dataset used for this analysis and its associated metadata are provided as a phyloseq object in Supplementary Dataset 2.

#### Domestic dog gut microbiome dataset

The dog analysis used 16S rRNA gene profiles from a globally distributed cohort of 261 domestic dogs sampled in Nepal, Thailand, the United Arab Emirates, and the United States across forager, agrarian, pastoralist, urban, and industrialized lifestyles (16). Four dogs with recent antibiotic exposure were excluded, leaving 257 dogs for the present analysis. The primary Archetypal Analyses used K = 8. Lifestyle was treated as an integrated ecological category reflecting correlated differences in human management, diet, environmental exposure, geography, climate, and degree of industrialization rather than as an isolated causal exposure. The exact dataset used for this analysis and its associated metadata are provided as a phyloseq object in Supplementary Dataset 3.

#### Visualization of archetype coefficients and taxonomic profiles

For each fitted model, sample-level archetype coefficients were displayed in two complementary ways. First, coefficients were mapped to a two-dimensional polygonal representation in which each archetype was assigned to a vertex and each sample position was calculated as the coefficient-weighted combination of the vertex coordinates. This display is an exact simplex representation only when K=3; for K>3, it is a two-dimensional barycentric projection of the K-component coefficient vector. Second, stacked coefficient plots displayed the full K-component mixture for each sample and therefore preserved the fitted coefficient values. Archetypal taxonomic profiles were represented in genus space. For the composition heatmaps, the five taxa with the largest contributions to each archetype were selected, pooled across archetypes, and deduplicated before visualization.

#### Bray-Curtis principal coordinates analysis

Conventional ordination was performed using Bray-Curtis dissimilarities calculated from the same samples and genus-level feature tables used for the corresponding Archetypal Analysis, following the same preprocessing steps. Principal coordinates analysis (PCoA) was applied to each Bray-Curtis distance matrix, and the first three principal coordinates were used for visualization.

For comparison with the sample distributions, archetypal profiles were projected into the fitted sample PCoA space as supplementary points using classical multidimensional-scaling interpolation based on their pairwise distances to the observed samples. This procedure placed the archetypes into the existing ordination without altering the sample coordinates.

#### Evaluation of K and sensitivity to model resolution

AA was evaluated across multiple candidate values of K. Diagnostic plots evaluated K = 3-10 for all three datasets. Biological metadata were not used to fit AA. For silhouette and Davies-Bouldin calculations, each sample was provisionally assigned to the archetype with its largest mixture coefficient. Higher silhouette values and lower Davies-Bouldin values indicate greater separation under this temporary hard assignment. Because AA represents continuous mixtures rather than discrete clusters, these statistics were treated as secondary diagnostics rather than as optimization criteria defining a uniquely optimal K.

Primary values of K were treated as representative resolutions chosen for parsimony and interpretability of the mixture representation. Coefficient-space displays across K were used to assess whether the principal biological organization persisted as model resolution changed. K = 7, K = 6, and K = 8 were used for the primary sample-source, early-life, and dog-lifestyle representations, respectively. Sensitivity maps spanning K=3-10 are shown for the sample source, TEDDY, and dog datasets in Figs. S2, S6, and S10, respectively.

#### Initialization and resampling stability

Initialization stability was evaluated by refitting AA to 100% of each dataset using five random initializations while keeping K and preprocessing fixed. Sampling stability was evaluated by refitting AA to training subsets comprising 50%, 60%, 70%, 80%, and 90% of samples, with five resampling replicates at each fraction. Differences between fitted archetypal taxonomic profiles were quantified using normalized discrete 1-Wasserstein distance. Archetypes were matched between runs to account for label permutation using the Hungarian algorithm (Fig. S13). For the resampling analysis, archetypes from each subsampled fit were compared with those obtained from the corresponding full-data solution.

#### Held-out projection onto fixed archetypes

To evaluate whether training-derived archetypes could serve as a fixed reference system for new observations, archetypes were learned using training samples only and then held fixed. For each withheld sample, a new non-negative coefficient vector constrained to sum to one was estimated by minimizing reconstruction error relative to the fixed archetypal profiles; the archetypes themselves were not refit using held-out samples. Coefficients were estimated using simplex-constrained least squares until the residual sum of squares changed by less than 10^-8^ or a maximum of 2,000 iterations was reached.

Held-out projections were evaluated across training fractions from 10% to 90% using silhouette, Davies-Bouldin, and PERMANOVA pseudo-F statistics calculated in the projected coefficient space (Fig. S14).

#### Random Forest analysis of AA-prioritized taxa

To test whether taxa emphasized by the archetypal profiles retained information about sample metadata outside the AA fitting procedure, taxa were ranked according to their contributions to the reconstructed archetypal taxonomic profiles. Progressively larger sets of AA-prioritized taxa were used as input features for Random Forest models, with duplicate taxa prioritized by more than one archetype retained only once. For each AA-prioritized feature set, control models were trained using equally sized sets of taxa sampled randomly from the eligible feature table.

Random Forest classification was used for sample-source and dog-lifestyle outcomes and evaluated using macro-F1. Random Forest regression was used for age in the early-life dataset and evaluated using R^2^. Models were implemented using scikit-learn RandomForestClassifier or RandomForestRegressor with 300 estimators and evaluated using five-fold cross-validation. The feature rankings were derived from the primary K=7, K=6, and K=8 archetypal solutions for the sample-source, TEDDY, and dog datasets, respectively.

#### Statistical analysis

Biological metadata were used only for post hoc interpretation of the primary unsupervised AA representations or for explicitly supervised validation analyses; they were not used to fit the primary AA solutions. Silhouette and Davies-Bouldin statistics were used as descriptive diagnostics of provisional hard assignments derived from AA coefficients. PERMANOVA was used in the held-out projection analysis as described above. Random Forest performance was evaluated using macro-F1 for categorical outcomes and R^2^ for continuous age prediction.

## Ethics statement

This study reanalyzed previously generated or publicly available, de-identified microbiome data and did not recruit or sample new human participants or animals. Ethical approvals, informed-consent procedures, permits, and animal-welfare procedures for the original data collections are described in the corresponding source publications (5, 7, 16).

## Data and code availability

All datasets analyzed in this study were previously generated or publicly available. The human multi-biospecimen data and the early-life samples were obtained from the resource described by Abdill et al. (7). The exact processed-data and their source are included in the Supplementary Datasets. The dog 16S rRNA gene sequence data are available in the NCBI Sequence Read Archive under BioProject PRJNA1472931 (16). All author-generated code required to reproduce AA fitting, K-sensitivity analyses, stability analyses, held-out projection, Random Forest validation, and figures are publicly available at https://github.com/RV2005/Archetypal-Analysis-Microbiome.

## Acknowledgments

We thank the New York University Abu Dhabi High Performance Computing (HPC) facility and UC Santa Cruz Genomics Institute compute cluster for computational support. This work was supported by the NYU Abu Dhabi Faculty Research Award (AD318) and a grant from the Bill and Melinda Gates Foundation (INV-008686).

