## Supplementary Figures for "Archetypal analysis provides interpretable mixture representations of microbiome community structure"

**Short title:** Archetypal representations of microbiome structure

**Competing-interest statement:** The authors have declared that no competing interests exist.

**Author Contributions:** Conceptualization: Aashish R Jha; Methodology: Ray Verma, Alexander G. Ioannidis, Aashish R Jha; Software: Ray Verma, Alexander G. Ioannidis; Data curation: Ray Verma, Aashish R Jha; Formal analysis: Ray Verma, Ava Albert; Visualization: Ray Verma; Resources: Alexander G. Ioannidis, Aashish R Jha; Supervision: Alexander G. Ioannidis, Aashish R Jha; Project administration: Aashish R Jha; Writing - original draft: Ray Verma, Aashish R Jha; Writing - review & editing: Ray Verma, Ava Albert, Alexander G. Ioannidis, Aashish R Jha.

Keywords: archetypal analysis; microbiome; dimensionality reduction; ordination; ecological gradients; community states; compositional data

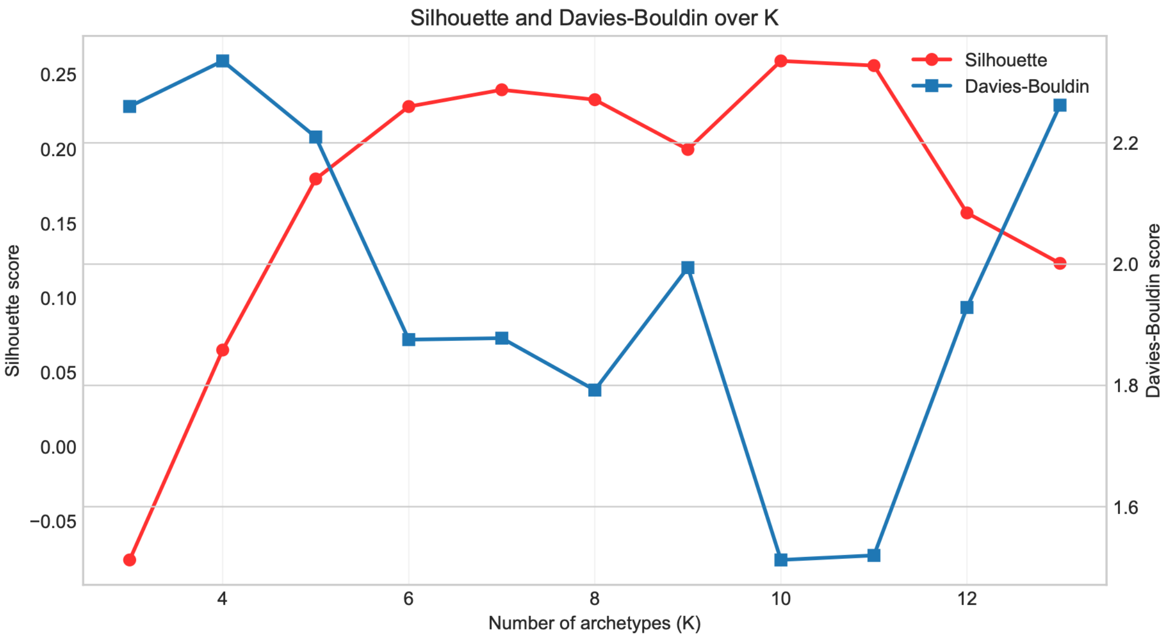


**Figure S1. Silhouette and Davies–Bouldin diagnostics across candidate values of K for the sample-source dataset.** Archetypal Analysis was fit across K=3–13, and each sample was provisionally assigned to the archetype with its largest mixture coefficient for calculation of silhouette and Davies–Bouldin statistics. Higher silhouette values and lower Davies–Bouldin values indicate greater separation under these temporary hard assignments. Because AA represents samples as continuous mixtures rather than discrete clusters, these statistics were treated as secondary diagnostics rather than as criteria defining a uniquely optimal K. Although K=10 produced the most favorable hard-assignment metrics within the tested range, K=7 was retained as a parsimonious intermediate resolution for the primary representation, with the major sample-source organization already evident at lower resolutions.

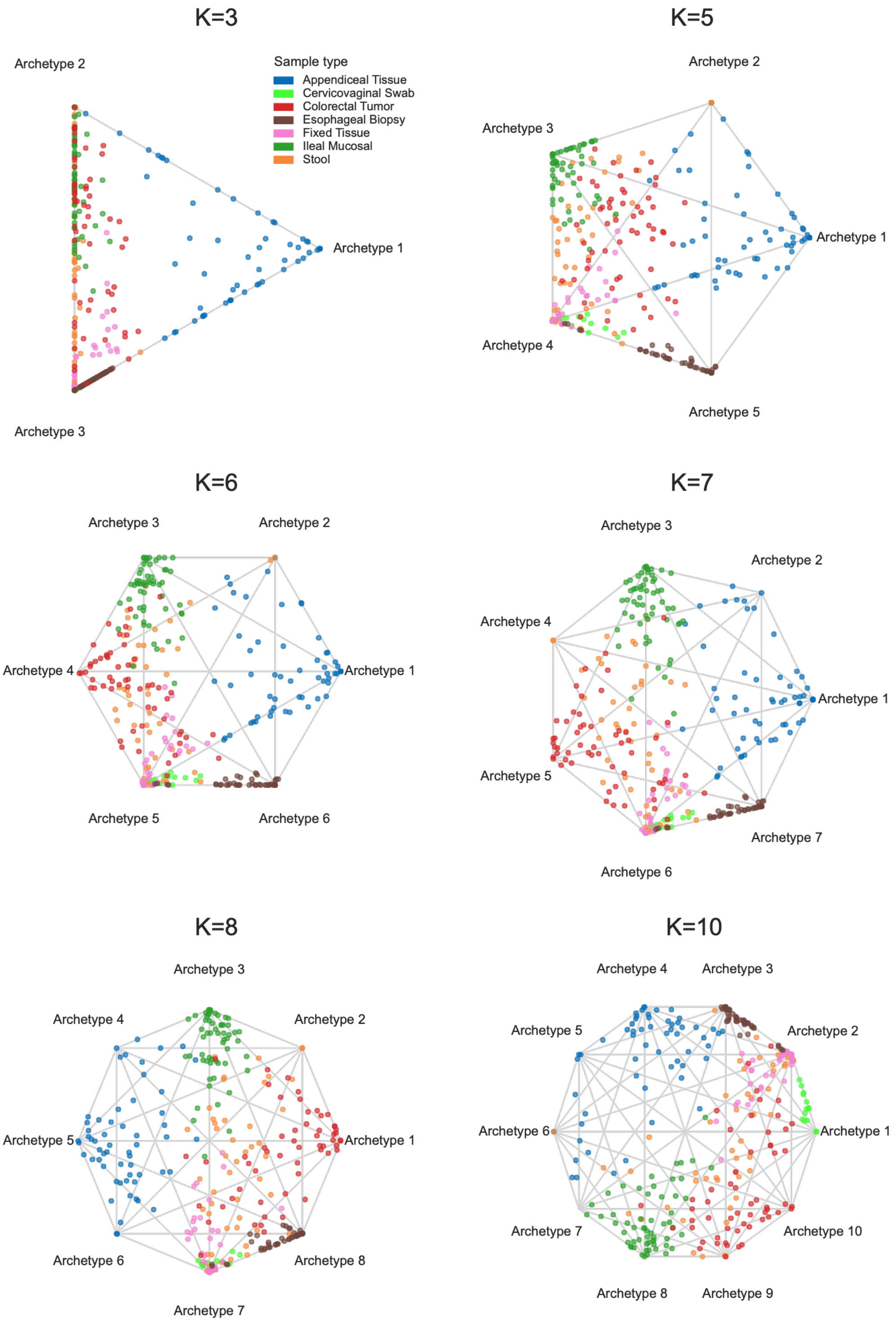


**Figure S2. Sensitivity of the sample-source archetypal representation to model resolution.** Archetypal representations of the multi-biospecimen dataset are shown for K=3, 5, 6, 7, 8, and 10. Samples are colored by sample-source category, with archetypes positioned at the vertices of the corresponding two-dimensional polygonal display. Across resolutions, the major sample-source organization remains recognizable, while increasing K allows additional archetypal structure to be represented. The primary analysis uses K=7; the broader series demonstrates that the principal biological interpretation is not dependent on a single exact value of K.

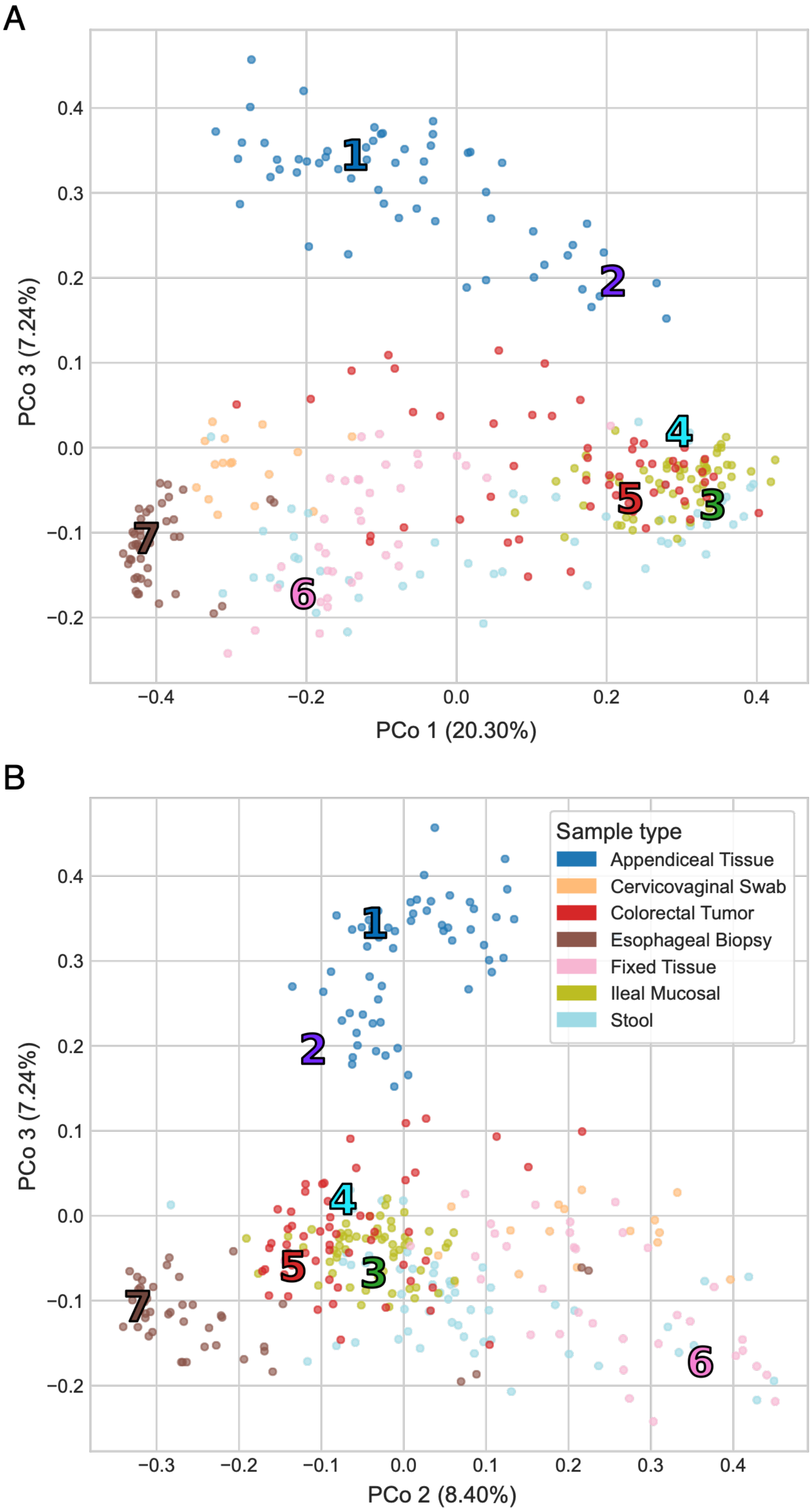


**Figure S3. Additional Bray–Curtis PCoA views of the multi-biospecimen microbiome dataset.** Bray–Curtis principal coordinates analysis of the same 319 samples shown in Fig. 2, with samples colored by sample-source category. **(A)** PCo1 versus PCo3. **(B)** PCo2 versus PCo3. Numbered markers indicate the locations of the seven archetypal profiles projected into the corresponding PCoA space. PCo1, PCo2, and PCo3 explain 20.30%, 8.40%, and 7.24% of the variation, respectively. These additional views illustrate sample-source structure not fully captured in the PCo1–PCo2 projection shown in the main figure.

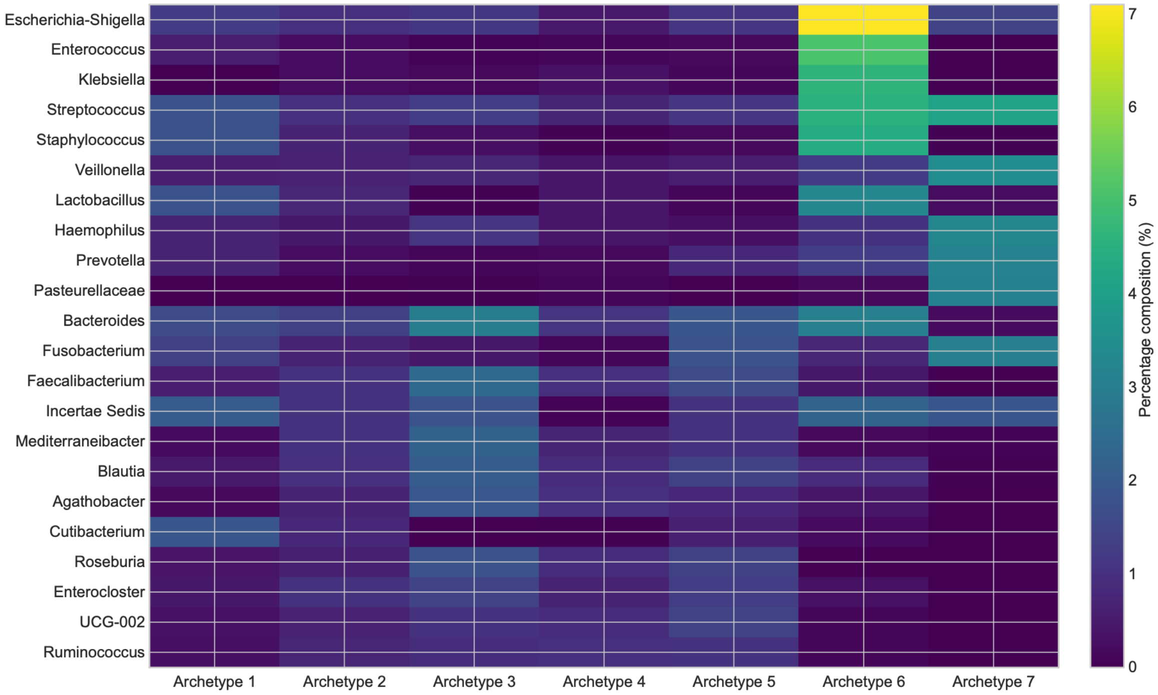


**Figure S4. Taxonomic composition of the sample-source archetypes.** Heatmap showing the taxonomic profiles of the seven archetypes in the multi-biospecimen dataset K=7. For visualization, the five taxa contributing most strongly to each archetype were selected, pooled across archetypes, and deduplicated. Columns represent archetypes and rows represent the selected taxa; color intensity indicates the percentage contribution of each taxon to the corresponding archetypal profile. The distinct profiles illustrate that the archetypes correspond to different microbial community compositions rather than solely to geometric endpoints in coefficient space.

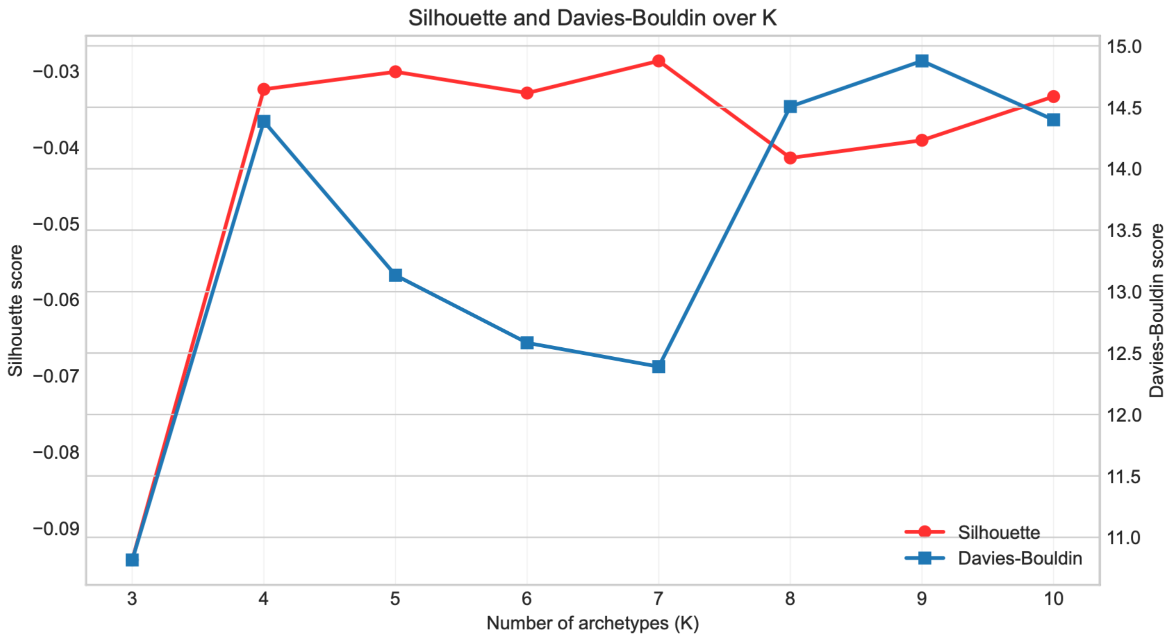


**Figure S5. Silhouette and Davies–Bouldin diagnostics across candidate values of K for the TEDDY early-life gut microbiome dataset.** Archetypal Analysis was fit across K=3–10, and each sample was provisionally assigned to the archetype with its largest mixture coefficient for calculation of silhouette and Davies–Bouldin statistics. Higher silhouette values and lower Davies–Bouldin values indicate greater separation under these temporary hard assignments. Because AA represents samples as continuous mixtures rather than discrete clusters, these statistics were treated as secondary diagnostics rather than as criteria defining a uniquely optimal K. The diagnostics improved markedly between K=3 and the intermediate resolutions but did not identify a single preferred value of K. K=6 was therefore retained as a parsimonious representative resolution, with sensitivity to alternative values evaluated separately in Fig. S6.

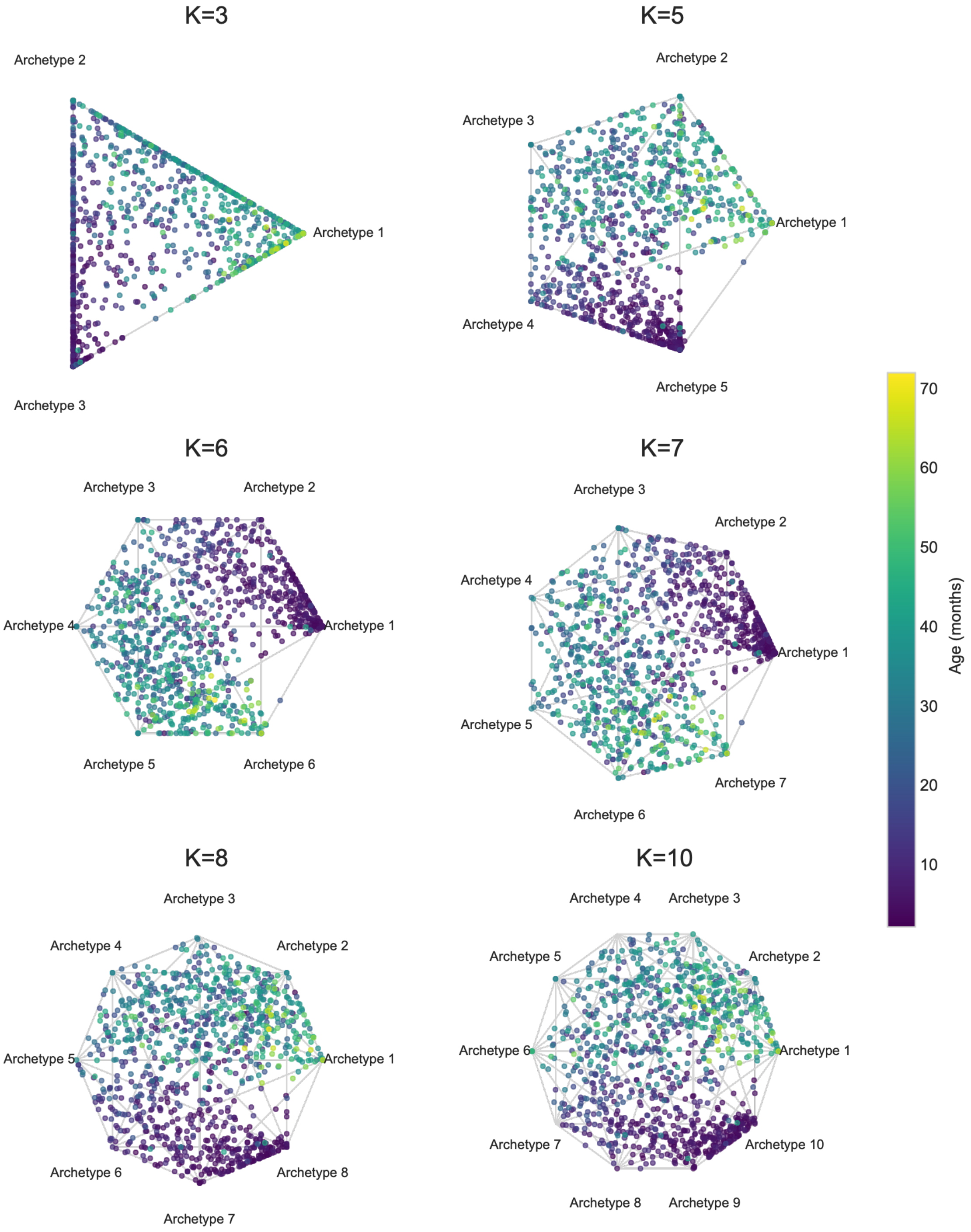


**Figure S6. Sensitivity of the early-life archetypal representation to model resolution.** Archetypal representations of the TEDDY early-life gut microbiome dataset are shown across selected values of (K). Samples are colored by age, and archetypes are positioned at the vertices of the corresponding polygonal display. Across resolutions, samples retain a graded age-associated organization, with younger and older microbiomes occupying different regions of archetype space and intermediate-age samples showing mixed archetypal contributions. The primary analysis uses K=6; the broader series demonstrates that the developmental organization is not dependent on a single exact value of K.

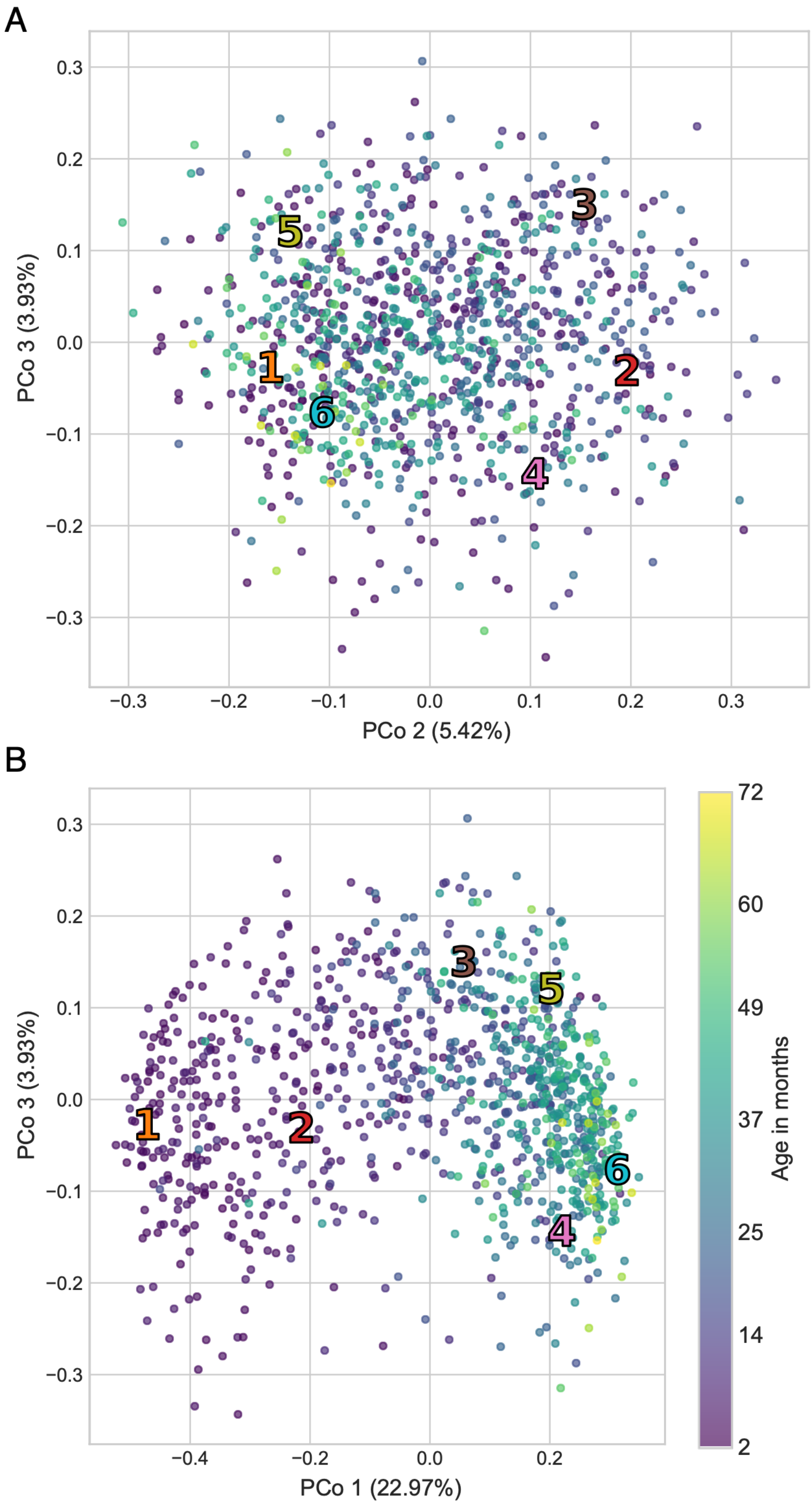


**Figure S7. Additional Bray–Curtis PCoA views of the TEDDY early-life gut microbiome dataset.** Bray–Curtis principal coordinates analysis of the same 1,033 samples shown in Fig. 3, with samples colored continuously by age in months. **(A)** PCo2 versus PCo3. **(B)** PCo1 versus PCo3. Numbered markers indicate the locations of the six archetypal profiles projected into the corresponding PCoA space. PCo1, PCo2, and PCo3 explain 22.97%, 5.42%, and 3.93% of the variation, respectively. These additional views complement the PCo1–PCo2 projection shown in the main figure and further illustrate the age-associated organization of the early-life gut microbiome.

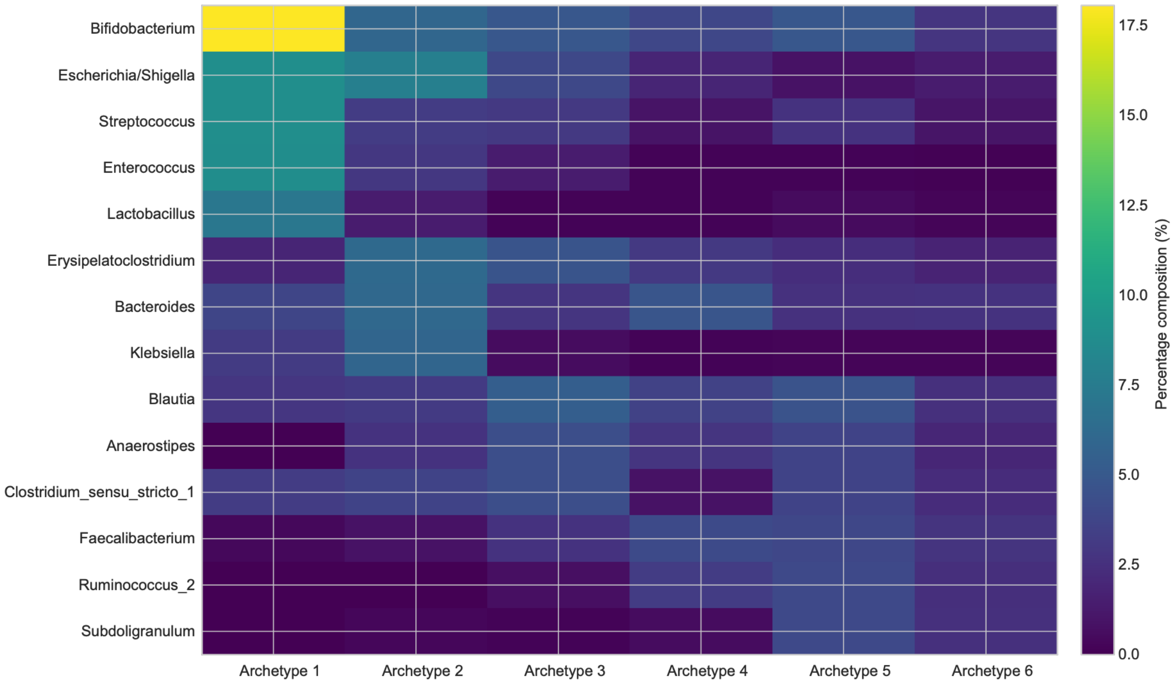


**Figure S8. Taxonomic composition of archetypes representing early-life gut microbiome succession.** Heatmap showing the taxonomic profiles of the six archetypes in the TEDDY early-life gut microbiome dataset K=6. For visualization, the taxa contributing most strongly to the archetypal profiles are shown; columns represent archetypes and rows represent taxa, with color intensity indicating percentage composition within each archetypal profile. Archetype 1, toward which the youngest samples are concentrated in Fig. 3, is characterized by a particularly strong contribution from *Bifidobacterium*, a hallmark genus of the early-life gut microbiome.

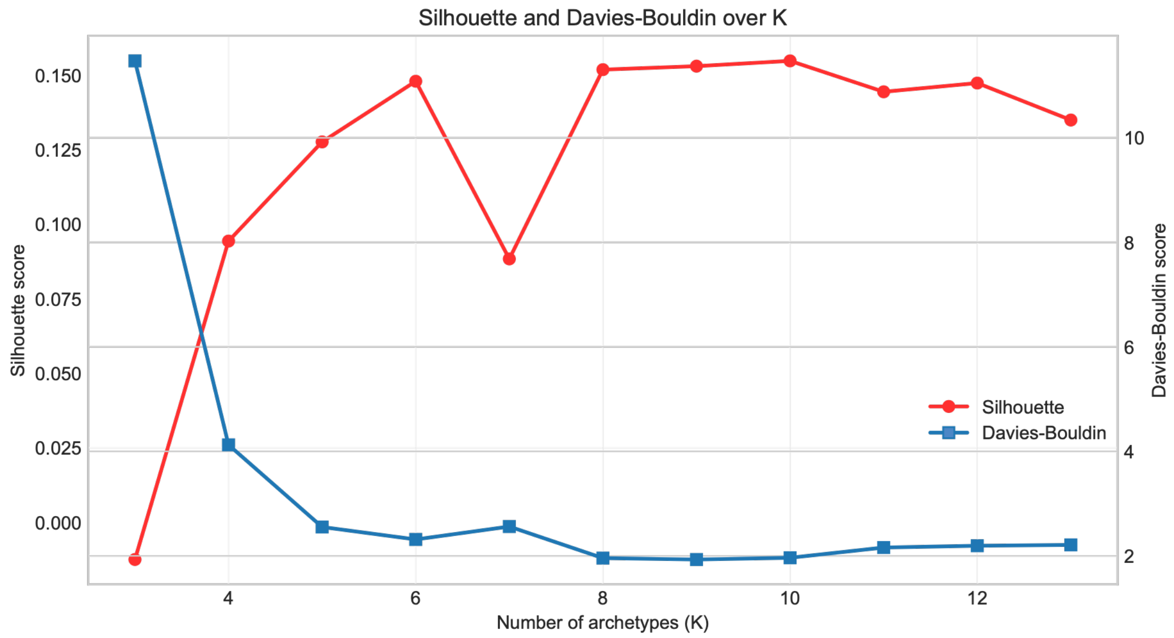


**Figure S9. Silhouette and Davies–Bouldin diagnostics across candidate values of K for the dog gut microbiome dataset.** Archetypal Analysis was fit across the tested range of K, and each sample was provisionally assigned to the archetype with its largest mixture coefficient for calculation of silhouette and Davies–Bouldin statistics. Higher silhouette values and lower Davies–Bouldin values indicate greater separation under these temporary hard assignments. Because AA represents samples as continuous mixtures rather than discrete clusters, these statistics were treated as secondary diagnostics rather than as criteria defining a uniquely optimal biological number of archetypes. Both metrics were favorable across approximately K=8–10, with relatively little improvement at higher resolutions. K=8 was therefore retained as the lowest parsimonious representative resolution within this favorable range.

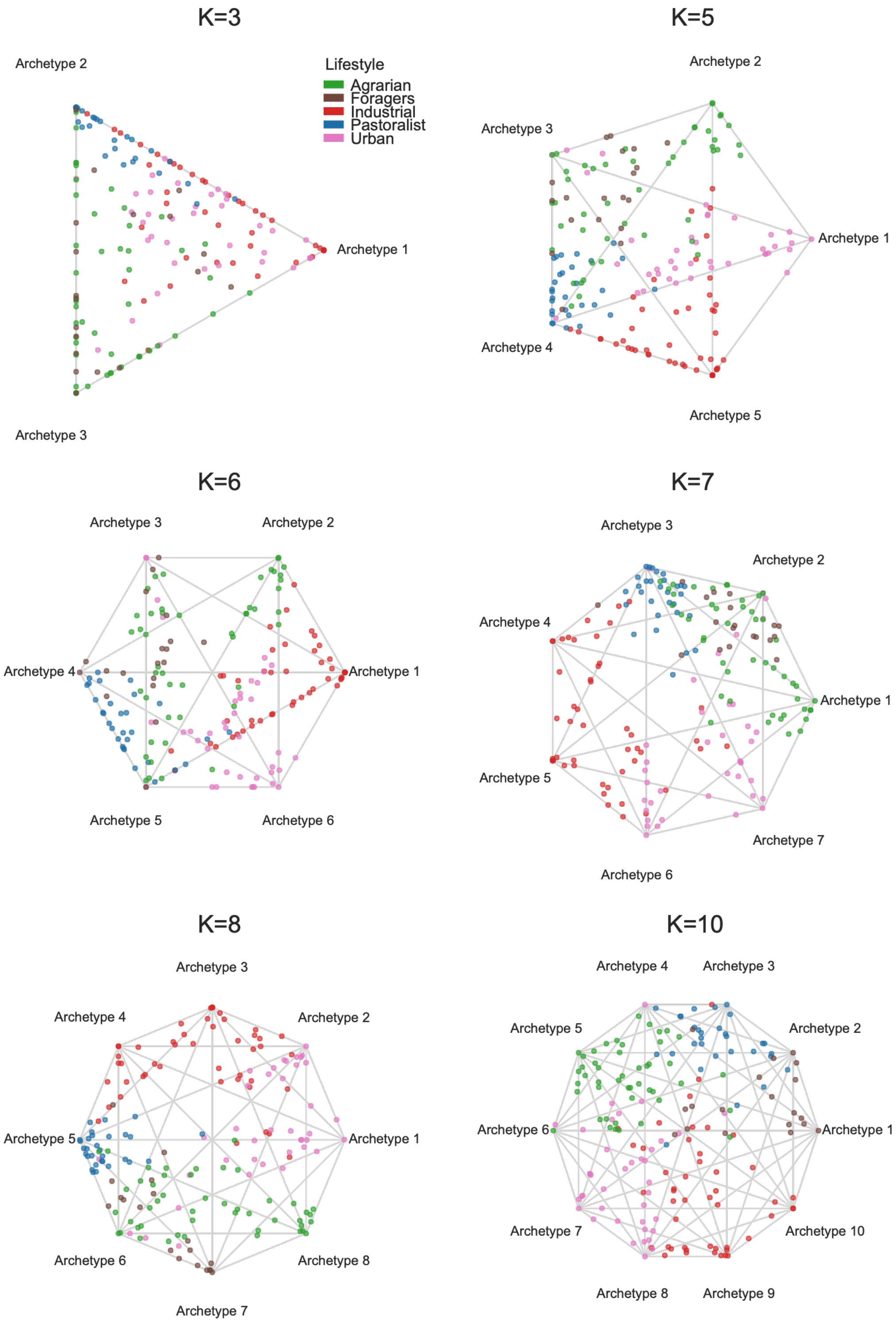


**Figure S10. Sensitivity of the dog microbiome archetypal representation to model resolution.** Archetypal representations of the dog gut microbiome dataset are shown for K=3, 5, 6, 7, 8, and 10. Samples are colored by lifestyle category, with archetypes positioned at the vertices of the corresponding two-dimensional polygonal display. Across resolutions, the major lifestyle-associated organization remains recognizable, while increasing K allows additional archetypal structure to be represented. The primary analysis uses K=8, the lowest representative resolution within the favorable range identified by the silhouette and Davies–Bouldin diagnostics shown in Fig. S9.

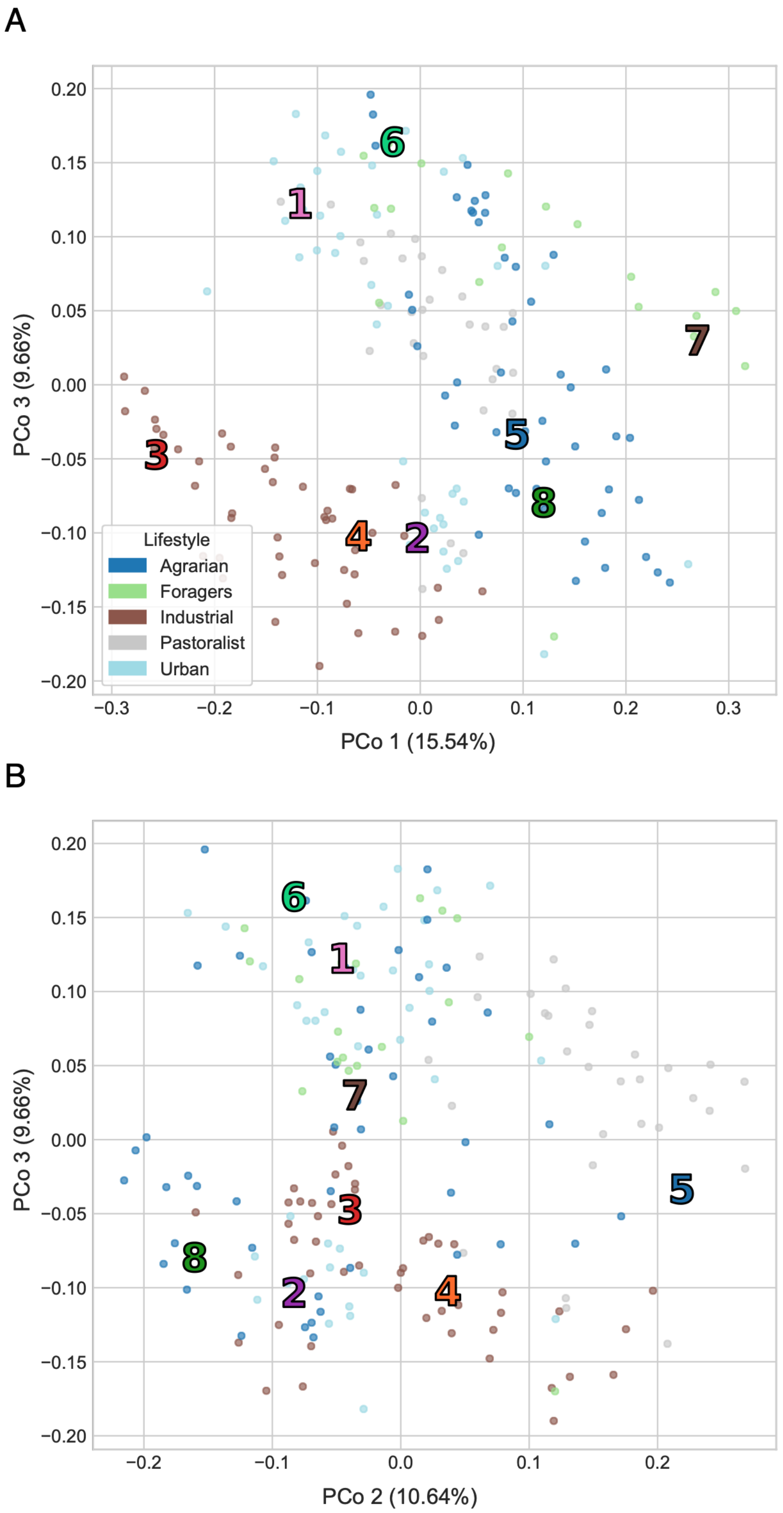


**Figure S11. Additional Bray–Curtis PCoA views of the dog gut microbiome dataset.** Bray–Curtis principal coordinates analysis of the same 257 dog microbiome samples shown in Fig. 4, with samples colored by lifestyle category. **(A)** PCo1 versus PCo3. **(B)** PCo2 versus PCo3. Numbered markers indicate the locations of the eight archetypal profiles projected into the corresponding PCoA space. PCo1, PCo2, and PCo3 explain 15.54%, 10.64%, and 9.66% of the variation, respectively. These additional views complement the PCo1–PCo2 projection shown in the main figure and further illustrate lifestyle-associated structure across the first three principal coordinates.

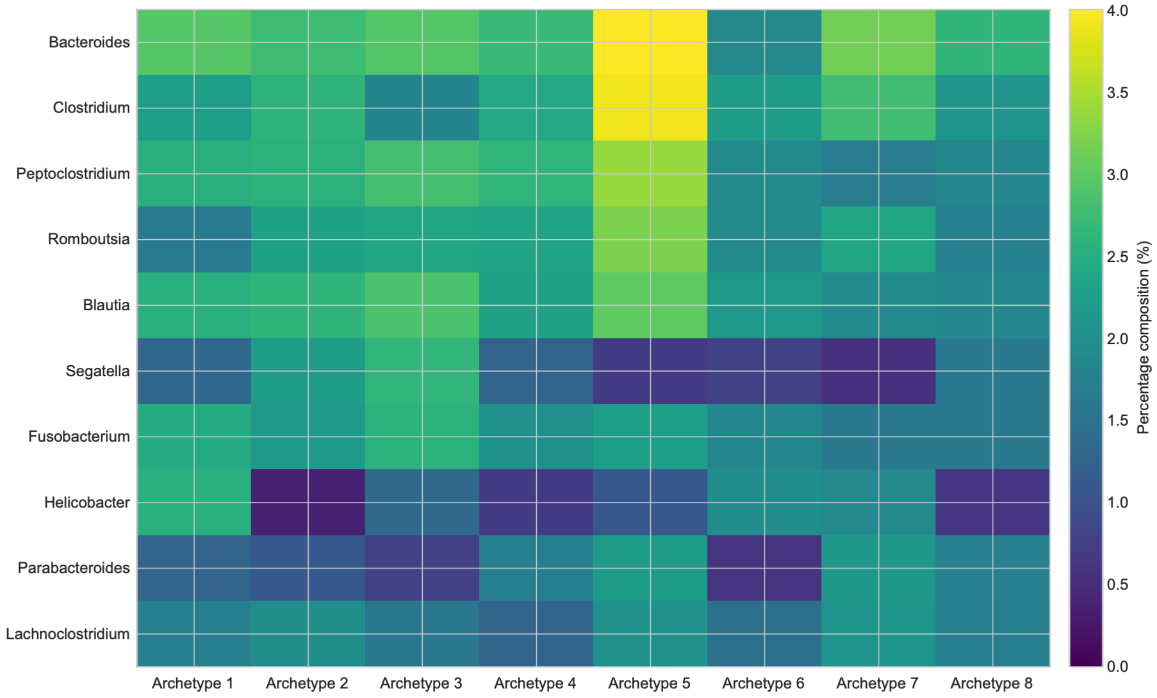


**Figure S12. Taxonomic composition of the dog gut microbiome archetypes.** Heatmap showing the taxonomic profiles of the eight archetypes in the dog gut microbiome dataset K=8. For visualization, taxa contributing most strongly to the archetypal profiles are shown; columns represent archetypes and rows represent taxa, with color intensity indicating percentage composition within each archetypal profile. The distinct taxonomic profiles illustrate that the archetypes represent different microbial community configurations underlying the lifestyle-associated structure shown in Fig. 4.

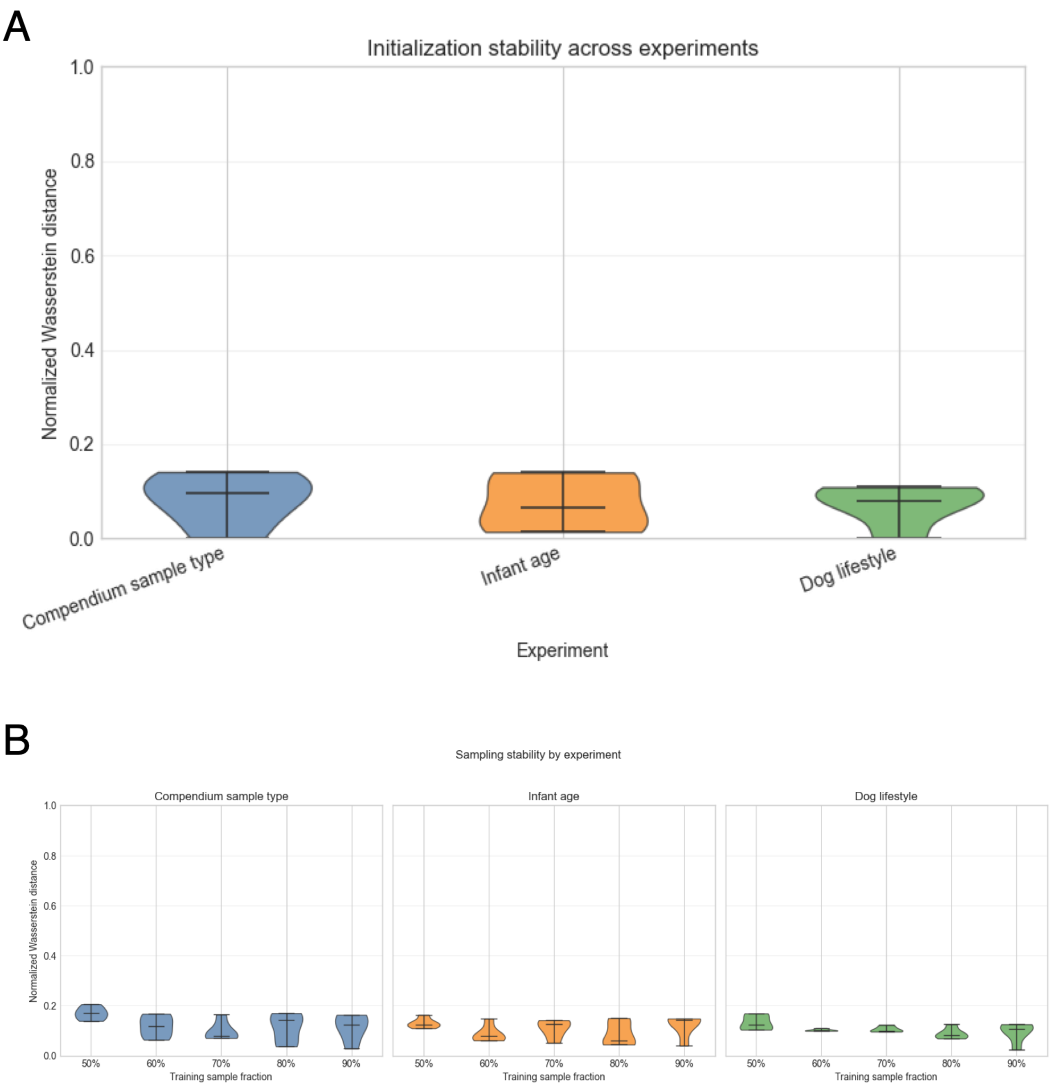


**Figure S13. Stability of archetypal profiles across initialization and data resampling. (A)** Initialization stability for the multi-biospecimen sample-source, TEDDY early-life, and dog-lifestyle datasets. Archetypal Analysis was refit using five random initializations for each dataset while holding K and preprocessing fixed. Archetypes were matched across runs after accounting for label permutation, and differences between matched solutions were quantified using normalized 1-Wasserstein distance, with lower values indicating greater agreement. Repeated initializations recovered broadly similar archetypal profiles across all three datasets. **(B)** Sampling stability across training subsets comprising 50%, 60%, 70%, 80%, and 90% of samples, with five random resampling replicates at each fraction. Archetypes from each subsampled fit were matched to those from the corresponding full-data solution and compared using normalized 1-Wasserstein distance. Archetypal profiles remained broadly similar across training fractions, although stability varied among individual resamples and datasets.

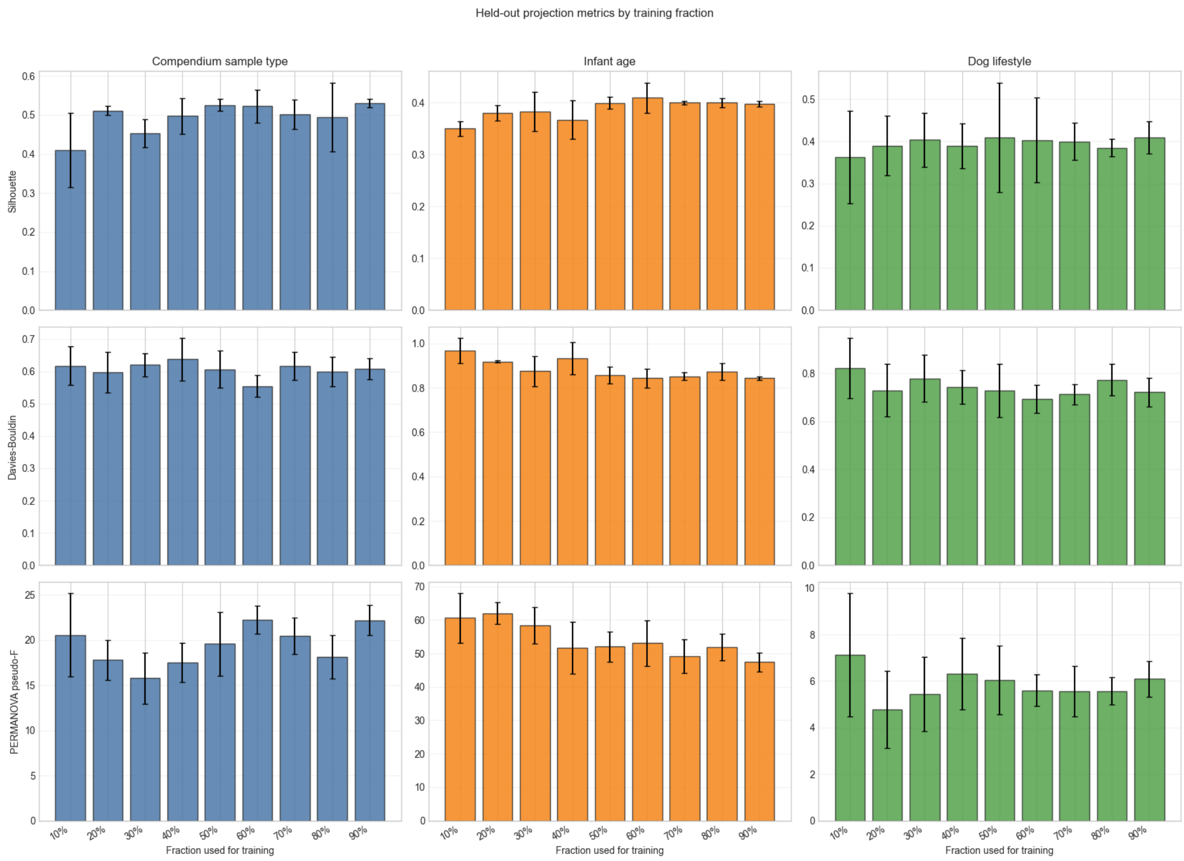


**Figure S14. Metadata-associated structure retained when held-out samples are projected onto archetypes learned from training data.** Archetypes were learned using training subsets comprising 10% to 90% of samples and then held fixed while the remaining samples were projected onto the learned archetypal representation. Columns show the multi-biospecimen sample-source, TEDDY early-life, and dog-lifestyle datasets, respectively. Preservation of metadata-associated organization among held-out samples was evaluated using silhouette score (top), Davies–Bouldin score (middle), and PERMANOVA pseudo-F statistic (bottom) in the projected coefficient space. Bars show results across training fractions, and error bars represent one standard deviation across repeated random splits. Across all three datasets, held-out samples retained recognizable metadata-associated organization over a broad range of training-set sizes, supporting the use of training-derived archetypes as a fixed reference system for new samples within the same dataset.
